# Dissecting the TMEM132A-EGFR Dependency to Unlock Translational Therapeutic Opportunities for Pan-Solid Tumor

**DOI:** 10.64898/2026.08.30.746586

**Authors:** Yaru Fu, Qiqi Ni, Can Ning, Junhao Wang, Xiang Fang, Mingyu Wu, Cheng Zhang, Jixian Wang, Jiayi Qian, Wentong Fang, Li Gong, Jing Yao, Dan Zhang, Xiaoming Li, Fei Zhao, Ninghong Song, Yuanqiao He, Xiyi Wei, Chao Qin, Jun Wang, Xijuan Liu

## Abstract

Solid tumors remain refractory to conventional treatments, yet cell surface proteins, by virtue of their extracellular accessibility and critical roles in tumor signaling, represent an attractive class of targets for precision-targeted therapy. Here, we report that transmembrane protein 132A (TMEM132A) is an essential and previously unrecognized pan-cancer target. TMEM132A interacts directly with EGFR and stabilizes its expression, thereby tethering EGFR at the plasma membrane and sustaining constitutive activation of lipid synthesis. Mechanistically, the TMEM132A-EGFR axis promotes lipogenesis by facilitating SREBP nuclear translocation, which in turn upregulates ACLY and ACSS2 expression to drive acetyl-CoA production and downstream lipid biosynthesis, ultimately disrupting lipid droplet homeostasis. To therapeutically target this axis, we developed a nanobody, LFNanoT132A#3, which effectively blocks the TMEM132A-EGFR interaction, abrogates downstream signaling activation, and potently inhibits proliferation across multiple solid tumor types. Our findings establish TMEM132A#3 as a critical node in membrane-tethered oncogenic signaling and metabolic rewiring, and position LFNanoT132A#3 as a promising therapeutic candidate for precision cancer therapy.

## Introduction

Solid tumors, originating in organs such as the lung, breast, and kidney, account for over 85% of all malignant cases and represent a leading cause of cancer-related mortality worldwide^1,2^. Unlike the remarkable efficacy of targeted therapies and immunotherapies in certain hematologic malignancies, solid tumors remain notoriously difficult to treat, largely due to their immunosuppressive microenvironment and genetic heterogeneity^3,4^. The precision oncology paradigm, exemplified by the success of EGFR inhibitors in lung cancer and HER2-targeted therapies in breast cancer, has revolutionized the treatment of molecularly defined tumor subsets^5^. However, not all solid tumors have benefited equally from this paradigm. ccRCC is driven by VHL loss-of-function in up to 90% of cases^6,7^, leading to HIF-mediated hypoxia signaling^8^, a mechanism that falls outside the oncogene-addiction framework targeted by current therapies. Triple-negative breast cancer (TNBC) similarly falls outside this paradigm due to the absence of ER, PR, and HER2 expression^9–11^. Even when targets exist, resistance is inevitable, as seen in lung adenocarcinoma, where EGFR inhibitors yield initial responses but acquired resistance almost universally emerges^12^. These challenges underscore an urgent, unmet need for novel therapeutic strategies that target common vulnerabilities across diverse solid tumor types.

Transmembrane proteins, owing to their unique biological localization and functional properties, represent promising targets for overcoming the therapeutic impasse in solid tumors. First, in terms of expression breadth, transmembrane proteins are commonly co-overexpressed or aberrantly activated across multiple solid tumor types, covering patient populations that are not adequately addressed by existing targeted strategies^13,14^. Second, with regard to functional significance, extensive studies have demonstrated that transmembrane proteins are broadly involved in key biological processes driving tumor initiation and progression, encompassing angiogenesis, energy metabolism, and immune regulation^15–17^. Third, in terms of druggability, the extracellular domains of transmembrane proteins are exposed on the cell surface, rendering them attractive targets for antibody-based therapeutics. Unlike nuclear transcription factors or cytoplasmic signaling molecules, transmembrane proteins can be recognized by drugs without traversing the cell membrane, thereby substantially reducing the delivery barrier and enhancing the feasibility of targeted therapy^13,18^.

Previous studies have established that TMEM132A activates the Wnt signaling pathway via interaction with the Wnt ligand transporter WLS in 293T cells^19^, TMEM132A also plays a critical role in embryogenesis^20^ and promotes prostate cancer growth under transcriptional regulation by E2F1^21^. Moreover, pan-cancer bioinformatics analyses have revealed that TMEM132A is highly expressed across diverse malignancies and correlates with poor patient prognosis^22^. Despite these insights, the functional breadth and therapeutic potential of TMEM132A as a broad-spectrum oncogenic driver remain underexplored.

In this study, through a CRISPR-Cas9 functional screen in ccRCC, a prototypical solid tumor driven by VHL inactivation and refractory to conventional therapies, followed by systematic pan-cancer validation, we identified TMEM132A as an essential oncogenic dependency across multiple tumor types. Mechanistically, TMEM132A sustains tumor growth by maintaining EGFR signaling and metabolic reprogramming. Notably, the extracellular nature of its functional interface allowed us to develop a specific blocking nanobody that exerts potent antitumor activity in diverse preclinical models. Our findings establish TMEM132A as a promising therapeutic target and provide a viable strategy for targeting a broad range of solid tumors.

## Result

### TMEM132A Loss Dampens ccRCC Tumorigenesis

We conducted a CRISPR-Cas9 screen using a Trafficking and Motility library in ccRCC cell line 786-O (Fig. S1A) and correlated the results with Hazard Ratio (HR) scores derived from TCGA data for ccRCC. To identify genes whose knockout impairs ccRCC cell proliferation, we selected genes that were significantly depleted in the negative selection screen. Among these, we further focused on genes with higher expression levels associated with increased HR, indicating that elevated expression predicts poorer prognosis in ccRCC patients. Candidate genes were ranked by HR values, and TMEM132A ranked 36th among the significantly depleted genes (Table S1, Fig.1A). Notably, TMEM132A is a direct target of SFMBT1 and ZHX2 (Fig. S1B), both of which are well-established key oncogenic drivers in ccRCC^23,24^, yet its own role in ccRCC remains unexplored. The convergence of prognostic significance, functional dependency in our screen, and regulation by known oncogenic transcription factors prompted us to prioritize TMEM132A for further investigation. By analyzing TCGA data, we found that higher stage (III and IV) tumors expressed higher levels of TMEM132A mRNA compared to lower stage (I and II) tumors and to normal tissue (Fig. 1B), and its high expression predicted worse prognosis in ccRCC (Fig. 1C). Next, to examine the physiological relevance of TMEM132A in ccRCC, we analyzed 9 pairs of tumor and normal tissues from ccRCC patients. Most tumors displayed (6 out of 9) consistent obvious upregulation of TMEM132A (both mRNA and protein) compared to paired normal tissues (Fig. S1C-D). Furthermore, we performed mIHC (multiplex fluorescent immunohistochemistry) on a tissue microarray (TMA) containing samples from 55 ccRCC patients (Table S2). The results showed that TMEM132A was expressed at a higher level in higher-stage ccRCC (Fig. 1D-E). Collectively, these data indicate that TMEM132A is highly expressed in ccRCC and its expression is associated with a worse prognosis.

**Figure 1.**
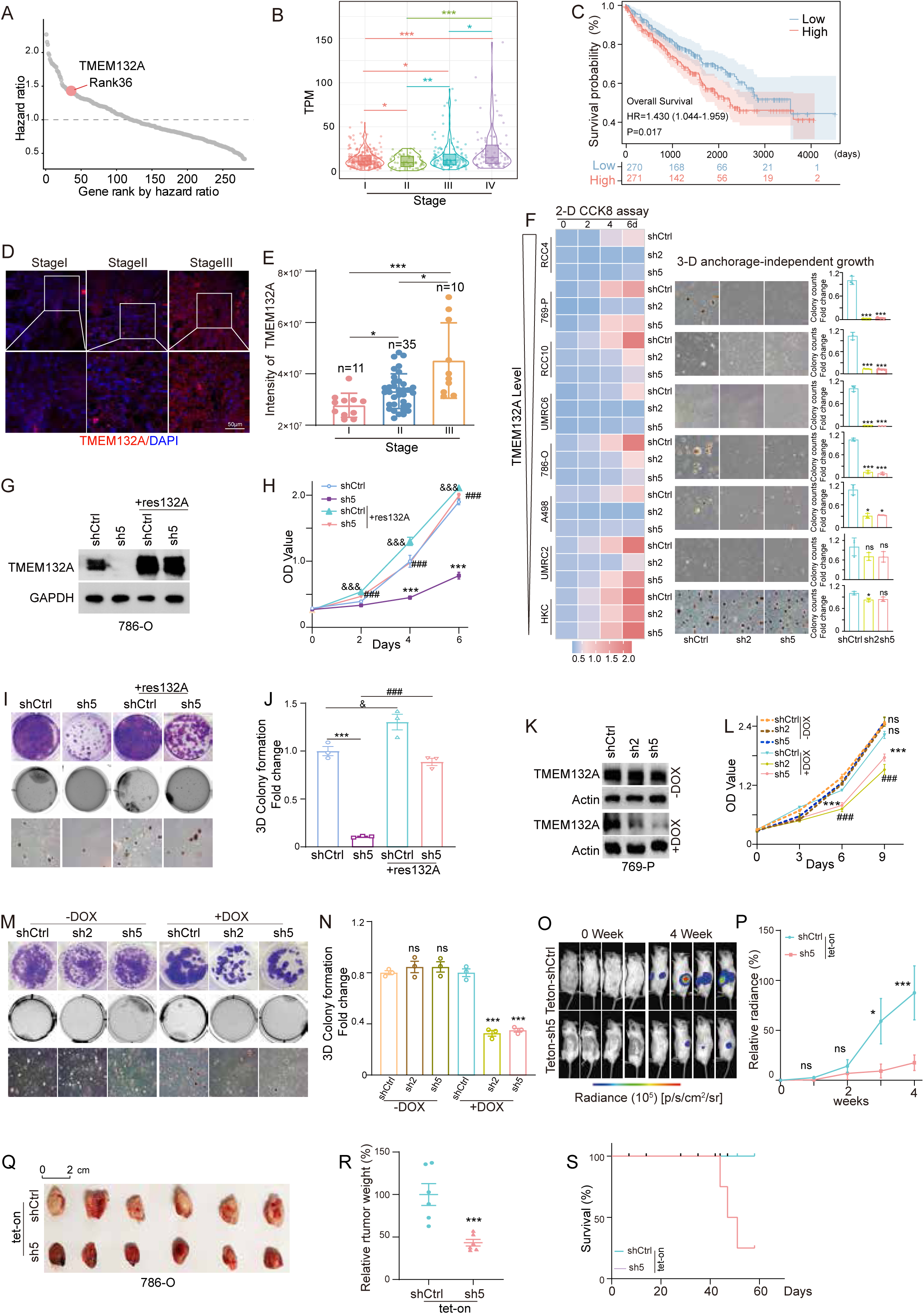
TMEM132A Loss Dampens ccRCC Tumorigenesis. (**A**) Distribution of genes ranked by the hazard ratio (HR) in the TCGA-KIRC cohort. (**B**) Violin plots showing TPM expression levels of TMEM132A in TCGA-KIRC tumors across stages I–IV. Each plot includes individual sample points with overlaid box plot summaries. Pairwise comparisons were performed using two-sided Welch’s t-tests, with statistical significance denoted by asterisks. (**C**) Kaplan-Meier plots indicating overall survival probability between high-and low-TMEM132A ccRCC patients. HR and P values are shown. Shaded areas indicate confidence intervals, and numbers below the plot show patients at risk. (**D**) Representative mIHC staining of TMEM132A in ccRCC patient tissues from stages I-III on TMA. (**E**) Quantification of TMEM132A staining. (**F**) Heatmap of CCK8 assays (left), representative images of 3-D soft agar assays (middle) and quantification of 3-D soft agar assays (right) in ccRCC and HKC cells transduced with either lentivirus expressing either shRNA control (shCtrl) or two individual TMEM132A shRNA (sh2 and sh5). (**G-J**) Immunoblots for lysates (**G**), CCK8 assays (**H**), colony formation assay (upper) and 3-D soft agar assays (lower) (**I**), and quantification of soft agar assays (**J**) of 786-O cells transduced with lentivirus encoding empty vector (EV) or TMEM132A-resistant (Res132A), followed by TMEM132A depletion. *sh5 versus shCtrl; ^#^sh5+Res132A versus sh5; ^&^shCtrl+Res132A versus shCtrl. (**K-N**) Immunoblots for lysates (**K**), CCK8 assays (**L**), colony formation assay (upper) and 3-D soft agar assays (lower) (**M**) and quantification of soft agar assays (**N**) of 769-P cells with TMEM132A depletion with or without doxycycline (DOX) treatment. *sh5 versus shCtrl; ^#^sh2 versus shCtrl. (**O-S**) Representative bioluminescence imaging (**O**), quantification of bioluminescence signals (**P**), tumor images (**Q**), tumor weights (**R**) and survival curves (**S**) post-DOX treatment after orthotopically injection of stable luciferase-expressing 786-O cells transduced with lentivirus expressing either inducible shCtrl and TMEM132A sh5. Data are shown as mean ± SEM, Statistical significance was determined by one-way ANOVA (**F**, **J** and **N**), two-way ANOVA (**H, L** and **P**) and Mann-Whitney test (**E** and **R**).

Having established that TMEM132A is highly expressed in ccRCC and correlates with poor prognosis, we next sought to determine its functional relevance in this malignancy. To this end, we examined TMEM132A levels in a panel of human ccRCC cell lines and renal proximal convoluted tubule epithelial cells (HKC). In line with the observations in patient tumors, TMEM132A mRNA and protein levels were increased in most of the ccRCC cells (Fig. S1E). To investigate the functional roles of TMEM132A in ccRCC, we depleted its expression using two independently validated shRNAs (sh2 and sh5) delivered in a PLKO-based lentiviral vector. Depletion of TMEM132A significantly decreased cell proliferation and 3D soft-agar growth in ccRCC models, and this effect was correlated with TMEM132A expression levels (Fig. 1F, Fig.S1F-G). In contrast, no such effect was observed in HKC cells, which exhibit low baseline levels of TMEM132A (Fig. 1F, Fig. S1F-G). To further confirm that this phenotype was due to the on-target effects of TMEM132A hairpins, we overexpressed (OE) shRNA-resistant TMEM132A in ccRCC cells and then depleted TMEM132A expression (Fig. 1G, Fig. S1H). Whereas the TMEM132A shRNA decreased cell proliferation and colony formation, shRNA-resistant TMEM132A OE efficiently rescued the phenotype (Fig. 1H-J, Fig. S1I-K), suggesting that these phenotypes were due to the on-target consequences of TMEM132A depletion. Subsequently, we designed two independent sgRNAs to target TMEM132A using CRISPR-Cas9 mediated elimination and observed a consistent phenotype (Fig. S1L-O).

To examine whether TMEM132A was important for ccRCC tumor growth, we generated two inducible TMEM132A shRNAs, which efficiently depleted TMEM132A levels upon doxycycline (DOX) addition in ccRCC cell lines (Fig. 1K, Fig. S1P). TMEM132A depletion also showed decreased cell proliferation and soft-agar growth upon DOX addition in ccRCC cells (Fig. 1L-N, Fig. S1Q-S). Next, either control or TMEM132A sh5 cells were orthotopically injected into the renal capsules of NCG mice. Tumor-bearing mice were confirmed by weekly bioluminescence imaging prior to DOX administration, which induced shRNA expression for *in vivo* growth monitoring. While control hairpin-expressing cells showed robust proliferation within 4 weeks of DOX treatment, TMEM132A-targeting shRNA expression abrogated tumor growth (Fig. 1O–R) and significantly extended overall survival (Fig. 1S). Taken together, our results suggest that TMEM132A is important for ccRCC tumorigenesis both *in vitro* and in *vivo*.

### The Emerging Role of TMEM132A as a Pan-Cancer Therapeutic Target

TMEM132A was identified as a potential pan-cancer biomarker through computational analysis of several databases^22^. Having established the clinical relevance of TMEM132A in ccRCC, we next extended our investigation to assess its pan-cancer expression and functional significance across additional tumor types. We discovered that high expression of TMEM132A universally predicted worse prognosis across all cancer stages (I-IV) in the pan-cancer TCGA dataset (Fig. 2A). Notably, compared with the other TMEM132 proteins, TMEM132A showed high HR scores in most cancer types (Fig. S2A), indicating its specificity correlation with worse prognosis. Given these findings, we further investigated the pan-cancer role of TMEM132A. To validate the functional dependency across diverse tumor contexts, we examined eight cell lines spanning six cancer types with high HR scores for TMEM132A (Fig. S2A), and found that TMEM132A knockdown (KD) significantly decreased cell proliferation and 3D soft-agar growth (Fig. S2B-C).

**Figure 2.**
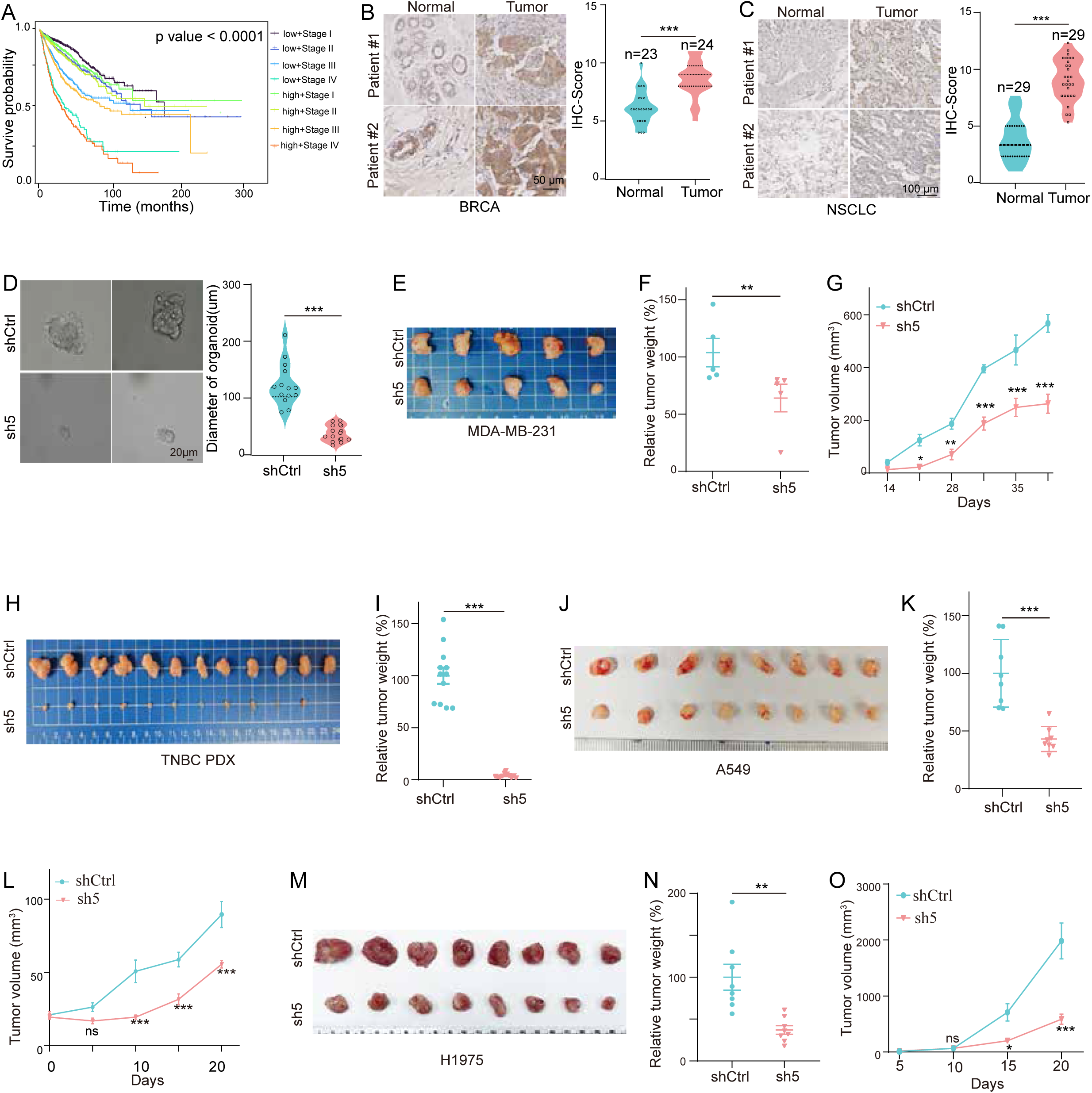
The Emerging Role of TMEM132A as a Pan-Cancer Therapeutic Target. (**A**) Kaplan-Meier overall survival curves for TCGA patients stratified by both TMEM132A expression level and pathological stage. The log-rank test showed significant survival differences among the combined expression-stage groups (P < 0.001). (**B-C**) Representative immunohistochemistry (IHC) staining images of TMEM132A in BRCA (**B**) or NSCLC (**C**) tumor tissue (T) and normal tissue adjacent tumor (N) (left); Quantification of TMEM132A staining (right). (**D**) Representative images (left) and quantification of diameters (right) of patient-derived breast cancer organoids in 3D culture transduced with TMEM132A depletion. (**E**-**G**) Tumor images (**E**), tumor weight (**F**) and tumor volume (**G**) from mice after orthotopically injection of MDA-MB-231 cells with TMEM132A depletion. (**H-I**) Tumor images (**H**) and tumor weight (**I**) from mice after orthotopically injection with TNBC PDX cells with TMEM132A depletion. (**J-L**) Tumor images (**J**), tumor weight (**K**) and tumor volume (**L**) from mice after subcutaneous injection of A549 cells with TMEM132A depletion. (**M-O**) Tumor images (**M**), tumor weight (**N**) and volume (**O**) from mice after subcutaneous injection of H1975 cells with TMEM132A depletion. Data show mean ± SEM. Statistical significance among groups was determined by Mann-Whitney test.

Considering the high HR of TMEM132A in BRCA and LUAD, we then focused on breast cancer (BRCA) and non-small cell lung cancer (NSCLC) as representative models. Immunohistochemical (IHC) staining revealed that TMEM132A expression was higher in these tumor tissues compared to their respective adjacent normal tissues (Fig. 2B-C). To model mammary carcinogenesis, we depleted TMEM132A in patient-derived mammary tumor organoids. Morphological and quantitative analyses revealed that TMEM132A depletion markedly impaired organoid growth (Fig. 2D, Fig.S2D). To assess the tumor-oncogenic role of TMEM132A *in vivo*, we established cell line-derived CDX models by orthotopically injecting MDA-MB-231 cells into the mammary fat pad, and found that TMEM132A depletion significantly reduced tumor growth, as evidenced by decreased tumor weight and volume (Fig. 2E-G). A comparable growth-inhibitory effect was further confirmed in a PDX model following TMEM132A KD (Fig. 2H-I, Fig. S2E). Similarly, depletion of TMEM132A in the murine breast cancer cell line EMT6 significantly inhibited cell growth (Fig. S2F-H).

Following KD of TMEM132A in A549 and H1975 cell lines, the cells were implanted subcutaneously into mice. Consistently, TMEM132A knockout markedly suppressed tumor growth in both CDX models (Fig. 2J-O). Notably, these two cell lines collectively recapitulate the major clinical challenge in NSCLC: A549 is intrinsically insensitive to TKIs, while H1975 harbors the acquired T790M resistance mutation. Taken together, these findings suggest that TMEM132A may represent a potential therapeutic target for NSCLC patients who are refractory or resistant to current EGFR-TKIs.

Collectively, our data demonstrate that TMEM132A plays a critical oncogenic role in cancers. Depletion of TMEM132A consistently suppresses tumor proliferation in diverse experimental settings, including organoids, cell line-and patient-derived xenografts. Notably, the efficacy of TMEM132A targeting in TKI-insensitive lung cancer cells highlights its potential as a compelling therapeutic target.

### TMEM132A Dysregulates Lipid Homeostasis across Multiple Cancer Types

To investigate how TMEM132A affects tumorigenesis, we performed RNA sequencing (RNA-seq) following the depletion of TMEM132A (Fig. 3SA). Gene Ontology (GO) analysis revealed that TMEM132A-regulated transcripts were enriched in genes associated with the Wnt signaling pathway (Fig. 3A), consistent with a prior study identifying TMEM132A as a novel regulator of Wnt signaling via interaction with WLS in 293T cells^19^. In addition, TMEM132A modulated the expression of genes involved in the metabolism of lipids including phospholipids, glycolipids, and fatty acids (Fig. 3A) -key constituents of lipid droplets (LDs)^25^. Subsequently, metabolomic profiling revealed a sharp decrease in acetyl-CoA levels upon TMEM132A depletion (Fig. 3B). As acetyl-CoA is the fundamental precursor for the synthesis of lipids^26^, we consistently observed a concomitant reduction in a spectrum of downstream metabolites, including steroids and sterols derivatives, glycerophospholipids, and fatty acids (Fig. S3B). Together, these results indicate that TMEM132A is a critical regulator of lipid metabolism, coordinating gene expression and metabolic flux to promote the synthesis of acetyl-CoA and its downstream lipid products. To further validate this finding, we employed a converse approach and found that TMEM132A OE significantly elevated acetyl-CoA levels (Fig. 3C). Accordingly, depletion of TMEM132A consistently reduced acetyl-CoA levels across multiple CDX models (Fig. 3D). To determine whether the acetyl-CoA deficit underlies the phenotypic effects of TMEM132A loss, we knocked down TMEM132A and treated several cancer cell lines with dichloroacetate (DCA), a PDK inhibitor that promotes pyruvate conversion to acetyl-CoA^27^. DCA treatment significantly rescued the TMEM132A-KD phenotypes, partially in 786-O and MDA-MB-231 cells and almost completely in A549 cells (Fig. 3E, Fig. S3C), supporting the conclusion that TMEM132A promotes tumorigenesis, at least in part, through maintaining acetyl-CoA levels and sustaining lipid metabolic programs.

**Figure 3.**
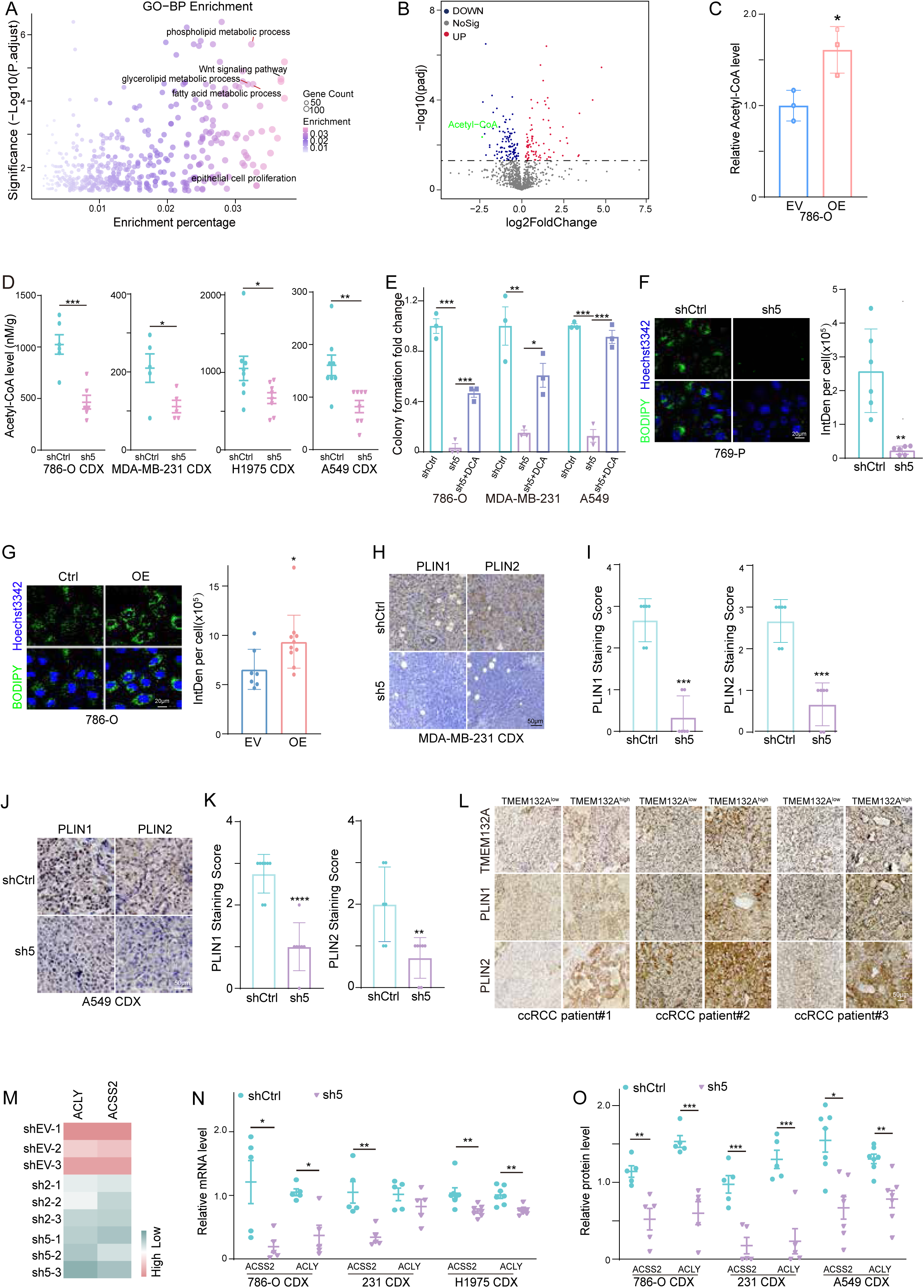
TMEM132A Dysregulates Lipid Homeostasis across Multiple Cancer Types. (**A**) GO biological process enrichment of DEGs (|Log2FC| > 1) from RNA-seq of 786-O cells between TMEM132A-sh5 and shCtrl. (**B**) Volcano plot showing differentially abundant (P<0.05) metabolites in 786-O cells (TMEM132A sh5 vs. shCtrl). (**C**) Relative acetyl-CoA level in 786-O cells infected with lentivirus encoding empty vector (EV) or FLAG-TMEM132A (OE). (**D**) Acetyl-coA levels in xenograft tumors harvested from mice injected with indicated cells. (**E**) Quantification of 3-D soft agar assays of indicated cell lines with TMEM132A depletion followed 250μM of DCA treatment. (**F**) Representative images (left) and quantitative analysis (right) of lipid droplet staining with BODIPY in 769-P cells with TMEM132A depletion. (**G**) Representative images (left) and quantification (right) of lipid droplet staining with BODIPY in 786-O cells transduced with lentivirus encoding EV or TMEM132A (OE). (**H-K**) Representative IHC staining (**H** and **J**) and dot plots showing quantification (**I and K**) of lipid droplet-associated protein PLIN1 and PLIN2 in xenograft tumors derived from orthotopically injected MDA-MB-231 cells and subcutaneously injected A549 cells. (**L)** Representative IHC staining of PLIN1 and PLIN2 in ccRCC patient tissues stratified by high (TMEM132A^high^) or low (TMEM132A^low^) TMEM132A expression. (**M**) Heatmap from RNA-seq analysis (n=3) of ACLY and ACSS2 expression profiles in 786-O cells with TMEM132A depletion. (**N-O**) qRT-PCR (**N**) and immunoblots quantification (**O**) of ACSS2 and ACLY in the indicated xenograft tumor tissues. Data are presented as means ± SEM. Statistical significance was determined by Student’s *t* test (**C**, **E**, **F** and **G**) and Mann-Whitney test (**D, I, K, N** and **O**).

Since phospholipids, glycerolipids, and fatty acids are key constituents of LDs, we hypothesized that TMEM132A contributes to LDs formation. To test this, we first assessed LDs formation using Bodipy staining in ccRCC cells following KD and OE of TMEM132A. We found that TMEM132A depletion consistently decreased LDs formation, while its overexpression increased it (Fig. 3F-G). Furthermore, in orthotopic TMEM132A-KD tumors, we observed a marked reduction in the levels of the lipid droplet-associated proteins perilipin 1 (PLIN1) and perilipin 2 (PLIN2) (Fig. 3H-K). To investigate the relationship between TMEM132A and LDs, we performed comparative analysis of PLIN1 and PLIN2 expression in representative ccRCC and BRCA tumor samples, focusing on regions with divergent TMEM132A expression (TMEM132A^high^ vs TMEM132A^low^). Notably, both PLIN1 and PLIN2 were consistently upregulated in TMEM132A^high^ regions and downregulated in TMEM132A^low^ regions (Fig. 3L, Fig. S3D). In summary, our findings show that TMEM132A perturbs the synthesis of acetyl-CoA and its downstream lipid products, ultimately disrupting LDs formation.

Next, we sought to determine how TMEM132A regulates acetyl-CoA levels. Given its established role in the transcriptional regulation of lipid metabolism (Fig. 3A), we interrogated our RNA-seq data. This revealed that TMEM132A depletion downregulated two critical enzymes in acetyl-CoA synthesis: ATP-citrate lyase (ACLY), which generates acetyl-CoA from citrate, and acetyl-CoA synthetase short-chain family member 2 (ACSS2), which converts acetate to acetyl-CoA (Fig. 3M). qPCR analysis further showed that the mRNA levels of ACLY and ACSS2 were restored by expression of shRNA-resistant TMEM132A, confirming the on-target effect (Fig. S3E). We therefore examined whether this transcriptional regulation was reflected *in vivo*. Indeed, tissues from the CDX model confirmed that TMEM132A KD robustly decreased both mRNA and protein levels of ACLY and ACSS2 (Fig. 3N-O, Fig. S3F). This coordinated downregulation at both transcriptional and translational levels strongly suggests that TMEM132A regulates acetyl-CoA synthesis through ACLY and ACSS2. To further validate the functional relevance of this pathway, we overexpressed TMEM132A and treated the cells with inhibitors targeting ACLY (ACLYi) or ACSS2 (ACSS2-IN-2). Each inhibitor alone effectively attenuated the TMEM132A-driven increase in cell proliferation, and their combined use yielded a mild synergistic suppression of cell growth (Fig. S3G-H). Taken together, our findings demonstrate that TMEM132A modulates acetyl-CoA production by transcriptionally regulating the key synthetases ACLY and ACSS2, thereby fueling lipid synthesis and promoting tumor growth.

### TMEM132A Interacts with EGFR to Promote Its Recycling

To investigate how TMEM132A promotes cancer cell proliferation, we performed FLAG Immunoprecipitation (IP) followed by mass spectrometry (IP-MS) in 786-O cells expressing either empty vector (EV) or FLAG-TMEM132A (TMEM132A OE) to identify the TMEM132A-binding proteins. Among the binding peptides, EGFR was one of the top hits in the TMEM132A OE (Fig. S4A), and given that TMEM132A KD was previously found to suppress NSCLC with wild-type and drug-resistant mutant EGFR (Fig. 2J-O), we selected EGFR as a candidate for further investigation. We undertook a multifaceted approach to validate the interaction between TMEM132A and EGFR. Initially, co-IP assays confirmed their binding in both exogenous and endogenous settings (Fig. 4A-B, Fig. S4B). We next employed immunofluorescence assay (IFA) in a panel of cancer cell lines, which showed predominant colocalization of FLAG-TMEM132A and endogenous EGFR at the cytoplasm and plasma membrane (Fig. 4C). To situate this interaction within a physiologic tissue architecture, mIHC analysis of BRCA organoids unequivocally demonstrated the colocalization of TMEM132A and EGFR (Fig. 4D). Collectively, these data conclusively demonstrate that TMEM132A interacts with EGFR and colocalizes *in vitro*.

**Figure 4.**
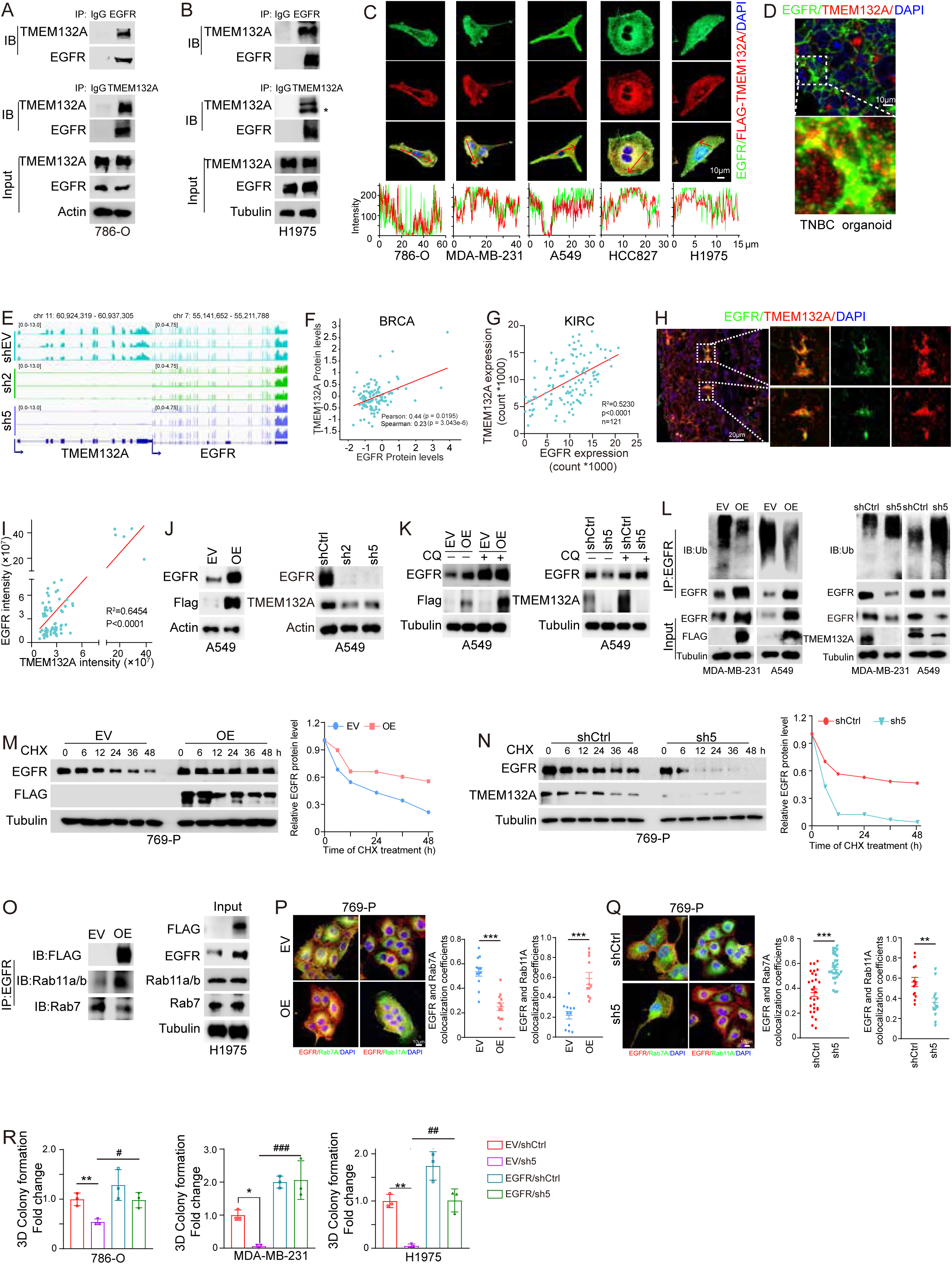
TMEM132A Interacts with EGFR to Promote Its Recycling. **(A**-**B**) Immunoprecipitation and immunoblots for lysates from 786-O cells (**A**) and H1975 cells (**B**). (**C**) Representative immunofluorescence assay (IFA) images (upper) in indicated cancer cell lines and quantification of the fluorescence intensity along the line embedded in the left images following the arrow direction (lower). (**D**) Representative mIHC co-staining images of TMEM132A and EGFR in TNBC organoid. (**E**) Genome browser tracks to show the RNA-seq signal of EGFR and TMEM132A in individual samples. RNA-seq analyses were based on three independent biological replicates. (**F)** Correlation between TMEM132A and EGFR protein levels in breast cancer patients. Each dot represents one patient. Pearson correlation statistics are shown as displayed by cBioPortal. (**G**) Correlation between TMEM132A and EGFR protein expressions in 121 monoclonal 786-O cells. Each dot represents one monoclonal. (**H-I**) Representative mIHC co-staining images (**H**) and quantification (**I**) of TMEM132A and EGFR in ccRCC patient tissues. (**J**) Immunoblots of EGFR and TMEM132A for cell lysates from A549 cells with TMEM132A OE (left) or depletion (right). (**K**) Immunoblots for cell lysates from A549 cells with TMEM132A OE (left) or depletion (right) followed 100μM Chloroquine (CQ) treatment overnight. (**L**) Ubiquitination level of endogenous EGFR in MDA-MB-231 and A549 cells following TMEM132A OE (left) or depletion (right). (**M-N**) Immunoblots for cell lysates from 769-P cells with TMEM132A OE (**M**) or depletion (**N**) followed 100μM cycloheximide (CHX) treatment at indicated intervals; The right panel shows the quantification of EGFR bands, with tubulin used as the loading control. EGFR expression levels in the EV/OE and shCtrl/sh5 groups at the 0-hour time point were normalized to 1.0. (**O**) Immunoprecipitation and immunoblots for lysates from H1975 cells with TMEM132A OE. (**P**-**Q**) Representative IFA images showing co-localization of EGFR with Rab7 or Rab11A in 769-P cells with TMEM132A OE (**P**) or depletion (**Q**). Quantitative analysis of co-localization was shown as Pearson’s coefficient. (**R**) Quantification of 3-D soft agar in indicated cell lines transduced with lentivirus encoding EV or EGFR followed by TMEM132A depletion. *EV/sh5 versus EV/shCtrl; ^#^ EGFR/sh5. versus EV/sh5. Error bars represent SEM. Statistical significance was determined by unpaired Student’s *t* test (**P, Q** and **R**).

We next interrogated our RNA-seq data and found that the mRNA level of EGFR remained unchanged upon depletion of TMEM132A (Fig. 4E). Furthermore, patient survival analysis revealed no significant differences among groups stratified by the mRNA levels of TMEM132A and EGFR, including those with high co-expression of both, high expression of either alone, or low expression of both (Fig. S4C), indicating that the co-expression of TMEM132A and EGFR transcripts lacks prognostic value and functional relevance at transcriptional level.

Notably, while tumors with EGFR gene amplification exhibited significantly higher average protein abundance compared to wild-type tumors (p= 4e^-16^), considerable overlap in protein abundance distributions between the two groups was observed. Moreover, the accuracy of predicting EGFR protein levels based solely on genomic amplification status was limited (average precision=0.37)^28^. These findings collectively indicate that EGFR regulation extends beyond genomic control. Given that TMEM132A physically interacts with EGFR, we therefore shifted our focus to post-transcriptional and protein-level regulatory mechanisms.

In contrast to the mRNA level, analysis of the cBioPortal database revealed a significant positive correlation between TMEM132A and EGFR protein levels in BRCA (Fig. 4F). Consistent with this, we observed a concordant relationship between TMEM132A and EGFR protein expression across 121 single-cell-derived 786-O colonies (Fig. 4G, Supplementary data, Table S3). Extending these observations to a physiological context, mIHC on ccRCC clinical samples demonstrated that the protein levels of TMEM132A and EGFR were spatially correlated, with TMEM132A^high^ regions expression coinciding with high EGFR expression, and vice versa (Fig. 4H-I). Collectively, these complementary approaches establish a consistent and significant positive correlation between TMEM132A and EGFR protein levels across multiple biological contexts. Thus, we sought to determine whether TMEM132A regulates EGFR abundance. Consistent with this hypothesis, gain-and loss-of-function experiments showed that EGFR protein levels were concomitantly increased by TMEM132A OE and conversely decreased upon TMEM132A depletion across a panel of pan-cancer cell lines (Fig. 4J, Fig. S4D). In addition, tissues from CDX models confirmed that TMEM132A KD significantly decreased EGFR protein levels (Fig. S3F). These data confirm that TMEM132A regulates EGFR at the protein level.

During EGFR trafficking, the receptor initially accumulates on the limiting membrane of early endosomes, where its fate is determined. EGFR that is sorted into the recycling pathway is recycled to the plasma membrane to sustain signaling, whereas EGFR that is targeted to late endosomes undergoes lysosomal degradation, resulting in signal termination^29^. Firstly, to determine whether TMEM132A regulates EGFR through lysosomal degradation, we employed the lysosome inhibitor chloroquine (CQ). CQ treatment effectively abolished TMEM132A-mediated regulation of EGFR across multiple cancer cell lines, as demonstrated in both gain-of-function and loss-of-function assays (Fig. 4K, S4E). Furthermore, ubiquitination assays revealed that TMEM132A OE suppressed EGFR degradation, whereas TMEM132A depletion promoted EGFR ubiquitin-mediated degradation (Fig. 4L, S4F). Consistent with these findings, TMEM132A OE significantly enhanced EGFR protein stability and prolonged its half-life, while TMEM132A depletion exerted the opposite effect (Fig. 4M–N, Fig. S4G–H). Our data collectively demonstrate that TMEM132A post-translationally regulates EGFR protein stability by modulating its lysosomal degradation.

To investigate how TMEM132A regulates the protein stability of EGFR, we considered the potential involvement of RAB GTPases, given that TMEM132A was found to interact with multiple Rab proteins in our IP-MS analysis (Fig. S4A). Among these, RAB5A is predominantly localized predominantly to early endosomes and initiates post-endocytic trafficking; RAB7A marks late endosomes and mediates cargo delivery to lysosomes for degradation; and RAB11A/B function as key regulators of the recycling pathway, facilitating the return of internalized proteins such as EGFR to the plasma membrane^29,30^. Strikingly, upon OE of TMEM132A, we observed enhanced interaction between EGFR and Rab11 but weakened association with Rab7 (Fig. 4O, S4I). Consistent with this, IFA revealed increased co-localization of EGFR with Rab11 and reduced overlap with Rab7, while TMEM132A knockdown yielded the opposite effects (Fig. 4P-Q, S4J-K). These findings suggest that TMEM132A promotes the rerouting of EGFR from the degradative pathway toward the recycling pathway, thereby enhancing its return to the cell surface and potentially increasing its stability. Mechanistically, we hypothesized that TMEM132A drives tumorigenesis by stabilizing EGFR. To test this, we examined whether restoring EGFR expression could rescue the tumor-suppressive effects of TMEM132A depletion. As expected, EGFR OE dramatically reversed the inhibition of tumor cell growth caused by TMEM132A loss (Fig. 4R, Fig. S4L-M).

In summary, our findings establish TMEM132A as a critical regulator of EGFR protein stability and tumorigenesis by reprogramming the endosomal trafficking of EGFR, favoring its Rab11-mediated recycling over Rab7-mediated degradation.

### The TMEM132A-EGFR Axis Promotes Lipogenesis by Promoting SREBP Nuclear Translocation to Enhance ACLY and ACSS2 Expression

KEGG analysis revealed that TMEM132A significantly modulated the PI3K-AKT, mTOR, and MAPK pathways (Fig. 5A). These pathways are canonical downstream effectors of EGFR signaling, which aligns with our earlier finding that TMEM132A interacts with EGFR. In light of the established role of the EGFR/PI3K/AKT axis in SREBP-dependent lipid homeostasis^31–33^, and because ACLY and ACSS2, both of which are regulated by TMEM132A (Fig. 3M), are downstream targets of SREBP1 and SREBP2, respectively, we next focused on SREBP. However, RNA-seq analysis revealed no significant changes in SREBP mRNA levels following TMEM132A depletion (Fig. S5A), thereby excluding a transcriptional mechanism.

**Figure 5.**
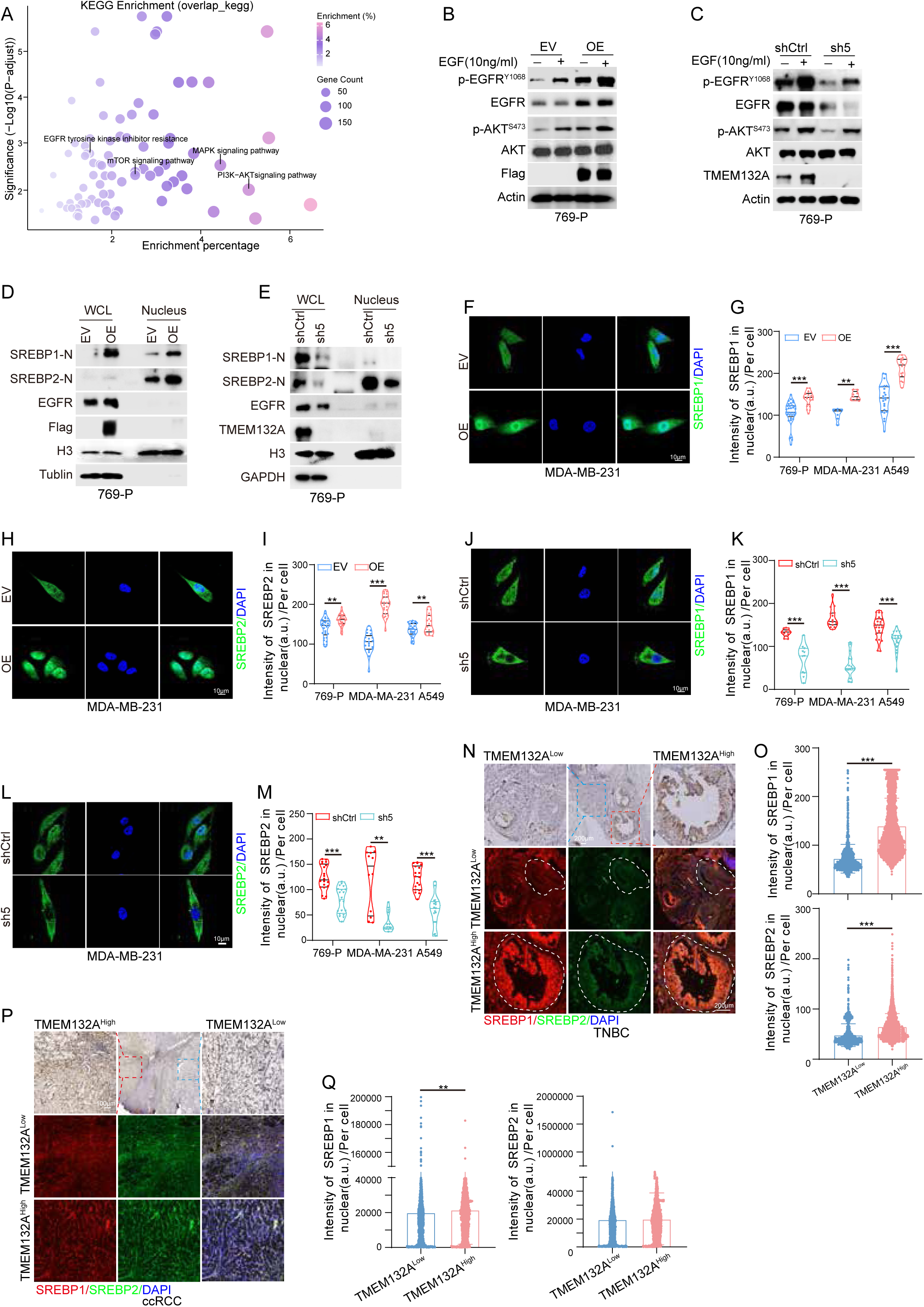
The TMEM132A-EGFR Axis Promotes Lipogenesis by Promoting SREBP Nuclear Translocation to Enhance ACLY and ACSS2 Expression. **(A)** KEGG biological process enrichment of DEGs from RNA-seq of 786-O cells between TMEM132A-sh2/sh5 overlapped genes and shCtrl. (**B-C**) Immunoblots for cell lysate of 769-P cells with TMEM132A OE (**B**) or depletion (**C**) followed with EGF (10 ng /ml) treatment for 1 h. (**D-E**) 769-P cells with TMEM132A OE (**D**) or depletion (**E**) were subjected to fractionation of whole cell lysates (WCL) and nucleus, followed by immunoblots analysis for the indicated proteins.(**F-M**) Representative IFA images of SREBP1 or SREBP2 in MDA-MB-231 cells with TMEM132A OE (**F** and **H**) or depletion (**J** and **L**); Quantification of the nuclear distribution intensities of SREBP1 (**G** and **K**) or SREBP2 (**I** and **M**) fluorescence intensity in indicated cell lines. (**N-Q**) Representative mIHC images (**N** and **P**) and quantification of the nuclear distribution intensities (**O and Q**) for SREBP1 and SREBP2 in TMEM132A^high^ and TMEM132A^low^ regions of breast cancer patient tissues (**N-O**) or ccRCC patient tissues (**P-Q**). Error bars represent SEM. Statistical significance was determined by unpaired Student’s *t* test (**G**, **I**, **K** and **M**) and Mann-Whitney test (**O** and **Q**).

Beyond transcriptional regulation, SREBP activation is critically regulated at the post-translational level. SREBPs are initially synthesized as hairpin-anchored membrane proteins in the ER. Upon transport to the Golgi, they undergo sequential proteolytic cleavage by S1P and S2P, releasing a soluble N-terminal transcription factor domain. The active fragment then translocates into the nucleus, where it binds to sterol regulatory elements (SREs) and activates target gene transcription^32,34^. We therefore examined whether TMEM132A modulates SREBP activity, specifically its nuclear localization via the EGFR/PI3K/AKT pathway.

We found that OE of TMEM132A robustly activated the EGFR signaling pathway and effectively increased pEGFR and pAKT levels comparable to those induced by EGF stimulation in several cancer cell lines (Fig. 5B, Fig. S5B). Conversely, TMEM132A depletion suppressed the EGFR pathway, and EGF failed to fully restore signaling to control levels (Fig. 5C, Fig. S5C). Taken together, these findings indicate that TMEM132A is a key regulator upstream of the EGFR/PI3K/AKT signaling pathway. Critically, unlike physiological EGFR activation, which depends on ligand binding, TMEM132A stabilizes EGFR and sustains constitutive EGFR signaling even in the absence of EGF stimulation. The ligand-independent EGFR signaling activation by TMEM132A acts as a pivotal molecular switch in cancer cells that effectively “locks” the EGFR pathway into a persistently active state, which in turn perpetuates uncontrolled proliferation.

Subsequently, we examined the functional consequence of this regulation on SREBPs. As expected, our data demonstrated that TMEM132A promoted nuclear SREBP (nSREBP), facilitating the entry of its transcriptionally active form into the nucleus (Fig. 5D-E, Fig. S5D-E). We also assessed SREBP nuclear translocation by IFA using gain-and loss-of-function models of TMEM132A. We observed a marked increase in SREBP nuclear intensity upon TMEM132A OE (Fig. 5F-I, Fig. S5F), and a corresponding decrease upon TMEM132A depletion (Fig. 5J-M, Fig. S5G). To assess the clinical relevance of TMEM132A and nSREBP, we performed mIHC staining on TNBC and ccRCC tissue samples stratified by TMEM132A expression levels. Regions with high TMEM132A expression exhibited stronger nuclear SREBP staining intensity (Fig. 5N-Q), although this association did not reach statistical significance for SREBP2 in ccRCC, likely reflecting intratumoral heterogeneity.

In conclusion, our findings delineate a coherent signaling axis wherein TMEM132A activates the EGFR/PI3K/AKT pathway, which in turn drives nuclear translocation of SREBP. This cascade ultimately leads to the transcriptional upregulation of key lipogenic enzymes ACLY and ACSS2, thereby promoting lipid metabolism reprogramming in cancer cells.

### Therapeutic Targeting of the TMEM132A–EGFR Interaction

To investigate the interaction interface between protein TMEM132A and EGFR, we employed AlphaFold3 to analyze their protein-protein interface, which identified two distinct sites of direct interaction (Fig. S6A). Subsequently, we constructed truncations of protein TMEM132A (aa392-635) and EGFR (aa25-645) that encompass these key regions. Pull-down assays demonstrated that the truncation of protein TMEM132A (aa392-635) binds full-length EGFR (Fig. S6B-6C), the truncation of EGFR (aa25-645) binds full-length protein TMEM132A (Fig. S6D), and the two truncated proteins also directly interact with each other (Fig. S6E).

To delineate the residues essential for TMEM132A-induced EGFR upregulation, we generated alanine-substitution mutants at sites predicted by AlphaFold3 and transiently expressed them in 293T cells. Notably, mutants W479A, R482A, R484A, and P497A lost the ability to upregulate EGFR expression (Fig. S6F). Next, we performed IP assays in 293T cells treated with CQ to block lysosomal degradation and stabilize protein levels. The results demonstrated that the W497A, R482A, R484A, and P497A mutations abrogated the interaction (Fig. S6G). Besides, these mutations also failed to upregulate EGFR in both 769-P and UMRC-2 cell lines (Fig. S6H). Phenotypically, W497A, R482A, R484A, and P497A mutations also impaired the ability of TMEM132A to promote ccRCC growth (Fig. S6I-J).

Since the interaction regions of TMEM132A and EGFR are both located in their extracellular domains, and the TMEM132A-EGFR interaction is essential for cancer cell proliferation, this prompted us to explore whether blocking this interaction could inhibit tumor cell growth. We therefore generated nanobodies using three peptides covering the aforementioned critical residues, yielding two candidates, LFNano1T32A#1 and #3 (Fig. 6A). Both nanobodies efficiently disrupted the TMEM132A–EGFR interaction, as demonstrated by semi-endogenous and endogenous IP assays (Fig. 6B–D). Notably, they also interfered with the binding between TMEM132A and the T790M-mutant EGFR in H1975 cells (Fig. 6D), underscoring their potential clinical relevance. Moreover, these nanobodies attenuated both EGFR upregulation induced by TMEM132A OE (Fig. 6E) and basal EGFR (Fig. 6F, left) or T790M-mutant EGFR (Fig. 6F, right) levels in cancer cells. Collectively, these findings indicate that the nanobodies are functionally effective in disrupting the TMEM132A–EGFR axis. Consistent with this, LFNanoT132A#1 and #3 suppressed proliferation of the ccRCC cell line 769-P (Fig. 6G) and abolished the growth advantage conferred by TMEM132A OE (Fig. 6H), confirming their on-target effects. Furthermore, the nanobodies also inhibited the growth of multiple other cancer cell lines, including neuroblastoma, prostate cancer, pancreatic adenocarcinoma, and lung cancer (Fig. S6K-N). Similar inhibitory effects were observed in BRCA PDX cell lines (Fig. 6I) and organoid models (Fig. 6J), further supporting the broad anti-tumor activity of these nanobodies. However, treatment with LFNanoT132A#1 similarly reduced the proliferation of HKC cells (Fig. 6K). Compared with LFNanoT132A#3, the impairing of LFNanoT132A#1 on non-malignant cells may warrant additional consideration when assessing its clinical potential.

**Figure 6.**
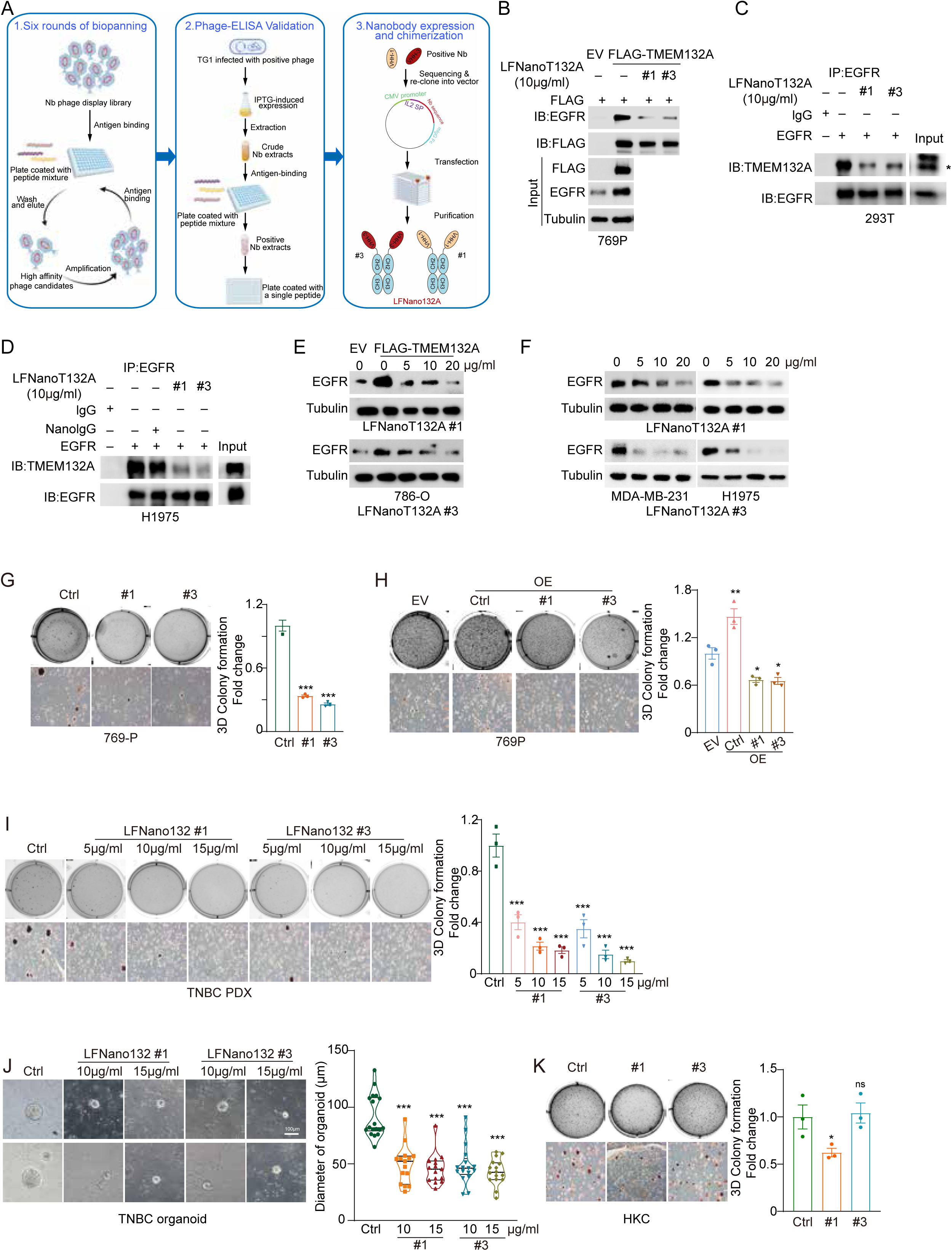
(**A**) Schematic overview of biopanning, ELISA screening, and chimerization of the nanobodies. (**B**) Immunoprecipitations and immunoblots for lysates from EV and TMEM132A OE 769-P cells after overnight incubation with 10μg/ml of LFNanoT132A#1 or #3 at 4°C. (**C-D**) Immunoprecipitations and immunoblots for lysates for 293T cell lysates (**C**) and H1975 cell lysates (**D**) after incubating with 10μg/ml of control nanobody (NanoIgG) or LFNanoT132A#1 and #3 overnight. (**E**) Immunoblots for lysates from EV and TMEM132A OE 786-O cells treated with indicated concentration of LFNanoT132A#1 (upper) or #3 (lower) for 72 h at 37°C. (**F**) Immunoblotting assay for lysates from MDA-MB-231 (left) and H1975 (right) cells treated with indicated concentration of LFNanoT132A#1 (upper) or #3 (lower) for 72 h at 37°C. (**G-H**) 3D soft agar assays (left) and quantification (right) of 3D soft agar assays of 769-P cells (**G**), EV and TMEM132A OE 769-P cells (**H**) with or without 10μg/ml of LFNanoT132A#1 or #3 treatment. (**I**) 3D soft agar assays (left) and quantification (right) of 3D soft agar assays of TNBC PDX cell line treated with indicated concentration of LFNanoT132A#1 or #3. (**J**) Representative images (left) and quantification of diameters (right) of patient-derived breast cancer organoids in 3D culture treated with indicated concentration of LFNanoT132A#1 or #3. (**K**) 3D soft agar assays (left) and quantification (right) of 3D soft agar assays of HKC cells treated with 10μg/ml of LFNanoT132A#1 or #3 treatment. Error bars represent SEM. Statistical significance was determined by one-way ANOVA.

### TMEM132A Nanoantibody (LFNanoT132A#3) Confers Broad-Span Therapeutic Vulnerability Against Multiple Solid Tumors

Next, we evaluated the antitumor efficacy of LFNanoT132A#1 and LFNanoT132A#3 in a murine 4T1 tumor model. To better recapitulate the complex tumor microenvironment, we selected immunocompetent Balb/c mice as the recipient model and orthotopically implanted tumor tissue cubes derived from donor mice into these recipients, rather than using conventional cell suspension inoculation, thereby preserving the native tumor architecture and stromal components. Treatments were then administered via intratumoral injection. Both antibodies effectively suppressed tumor growth (Fig. S7A–B) without inducing significant body weight changes (Fig. S7C). Notably, LFNanoT132A#3 exhibited a more significant difference in its antitumor efficacy compared to #1 (Fig. S7A). At endpoint necropsy, major organs (heart, liver, spleen and kidneys) showed no obvious histopathological abnormalities upon H&E staining across all groups (Fig. S7D). Considering both the anti-tumor efficacy and the detrimental impact of LFNanoT132A#1 on HKC cells (Fig. 6K), we selected LFNanoT132A#3 for further evaluation. The therapeutic potential of LFNanoT132A#3 was further substantiated in an additional syngeneic model, where it significantly suppressed Panc02 tumor growth in C57BL/6J mice (Fig. 7A–C). Notably, a progressive decline in body weight was observed in the control group during the later stages of the Panc02 study (Fig. 7D), likely attributable to excessive tumor burden, which necessitated premature termination of the experiment.

**Figure 7.**
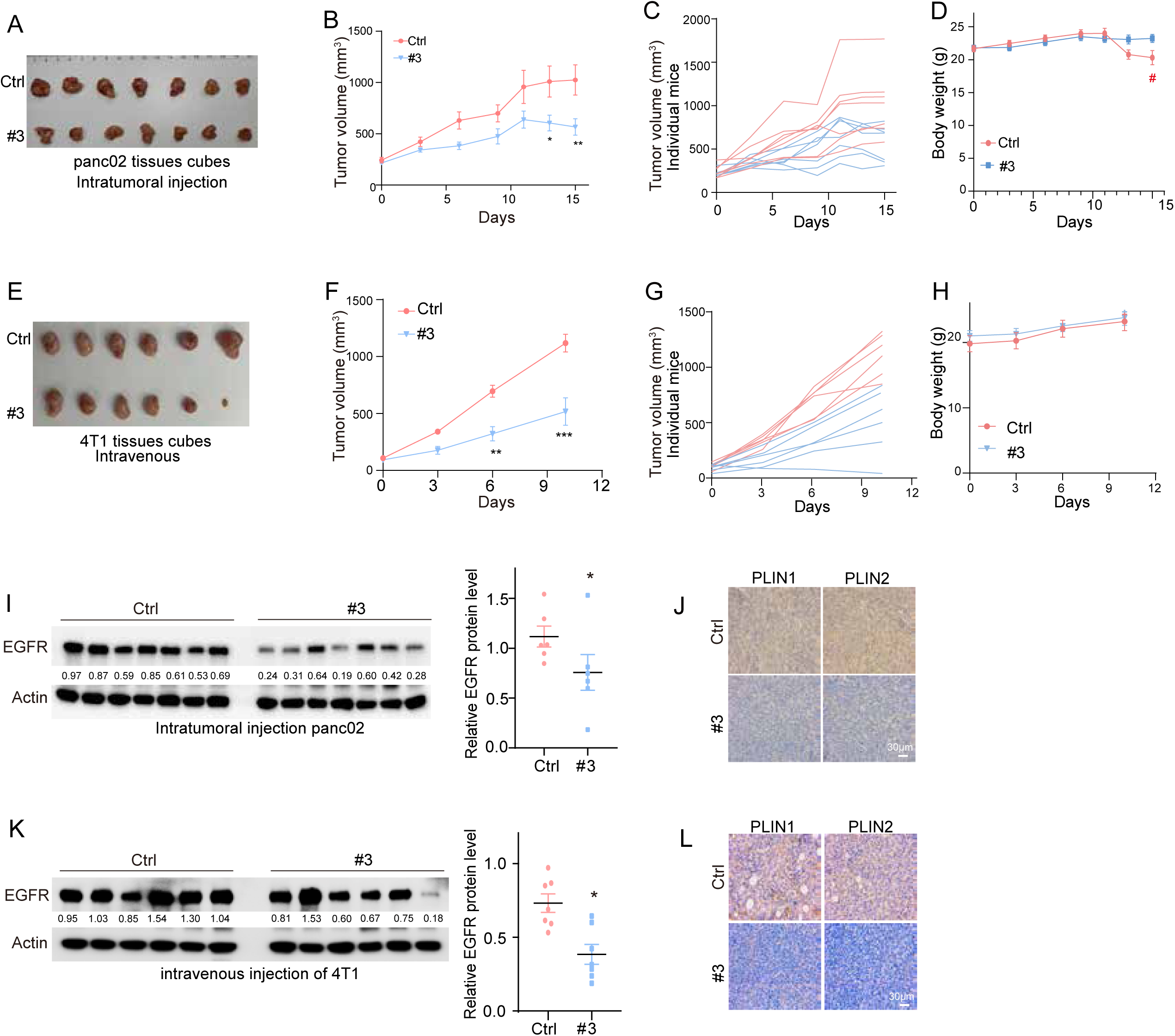
(**A-D**) Tumor images (**A**), tumor growth curve (**B**), individual tumor growth curves (**C**) and tumor weight (**D**) of C57BL/6J mice implanted with panc02 tumor tissue cubes and then intratumorally injected with LFNano132A #3. (**E-H**) Tumor images (**E**), tumor growth curve (**F**), individual tumor growth curves (**G**) and tumor weight (**H**) of Balb/c mice implanted with 4T1 tumor tissue cubes and then intravenously injected with LFNano132A #3. (**I**) Immunoblots for lysates from pan02 tumor tissues (left) and quantification of EGFR bands, with actin used as the loading control (right). (**J**) Representative IHC staining of PLIN1 and PLIN2 from pan02 tumor tissues. (**K**) Immunoblots for lysates from 4T1 tumor tissues (left) and quantification of EGFR bands, with actin used as the loading control (right). (**L**) Representative IHC staining of PLIN1 and PLIN2 from 4T1 tumor tissues. Data show mean ± SEM. Statistical significance among groups was determined by Mann-Whitney test.

To further assess the translational potential of LFNanoT132A#3, we also evaluated its activity via intravenous administration in the 4T1 model. Similar to the intratumoral route, intravenous delivery of LFNanoT132A#3 effectively inhibited tumor growth (Fig. 7E–G) without causing significant body weight loss (Fig. 7H), and histopathological analysis again showed no notable organ toxicity (Fig. S7E). Mechanistically, we assessed EGFR expression in tumor tissues following LFNanoT132A#3 treatment. A significant decrease in EGFR protein levels was observed (Fig. 7I and 7K), along with marked downregulation of PLIN1 and PLIN2 as determined by IHC (Fig. 7J and 7L). These data collectively indicate that LFNanoT132A#3 exerts its on-target activity in vivo, effectively disrupting the TMEM132A-EGFR axis and suppressing lipogenesis.

Collectively, these findings demonstrate that LFNanoT132A#3 exerts potent and broad-spectrum antitumor activity across multiple syngeneic models and administration routes, with a consistent safety profile characterized by the absence of overt toxicity. These results position LFNanoT132A#3 as a promising therapeutic candidate worthy of further preclinical development and translational investigation.

## Discussion

De novo lipogenesis is a core metabolic reprogramming event driving the progression of multiple solid tumors. Extensive studies have demonstrated that aberrant lipid metabolism is widely documented in hepatocellular carcinoma, breast cancer, colorectal cancer, lung cancer, and prostate cancer, and its aberrant activation is closely associated with tumor proliferation, invasion, metastasis, and poor prognosis^35,36^. Among these, ccRCC, characterized by abundant cytoplasmic lipid accumulation, has emerged as a representative model for studying lipid dependence in solid tumors. Lipid synthesis represents not only a shared metabolic vulnerability across multiple solid tumors but also a promising therapeutic target. In this study, we identified TMEM132A as a novel regulator of lipid synthesis that acts through EGFR modulation. Importantly, our findings further establish TMEM132A as a pan-cancer target, and we have delineated the mechanism by which it drives tumor progression through EGFR-dependent lipid synthesis.

Existing precision therapies, built upon molecularly defined tumor subsets, enable the selection of corresponding targeted agents through the identification of specific driver mutations such as EGFR. This approach shifts treatment from the "carpet-bombing" paradigm of conventional chemotherapy to a "precision-guided" strategy, substantially improving treatment response rates. However, many common solid tumors have not benefited equally from current precision therapies. ccRCC is driven by loss-of-function alterations that are intrinsically difficult to target^37^, whereas TNBC lacks high-frequency actionable driver mutations, leaving few targeted options for most patients^38^. Despite the clinical success of EGFR-TKIs in mutation-positive lung cancer, acquired resistance, notably via T790M, inevitably emerges. Beyond resistance, many EGFR wild-type patients do not respond to EGFR inhibitors and thus lack targeted treatment options^39^. These unmet clinical needs across diverse tumor types underscore the urgent demand for alternative therapeutic strategies that operate outside the conventional targetable-driver framework. Here, we turn to TMEM132A, a cell-surface protein, as a promising candidate to address this gap. Leveraging the unique advantages of TMEM132A as a cell-surface target, this study evaluates the oncogenic function of TMEM132A across multiple solid tumor models, including ccRCC, TNBC, and EGFR wild-type or T790M-mutant lung cancer. We aim to establish a promising therapeutic candidate and a mechanistic framework to accelerate its clinical translation.

In normal tissues, cellular proliferation is strictly governed by the regulated production and release of growth-promoting signals. Cancer cells, by contrast, acquire the hallmark ability to achieve self-sufficiency in growth signaling, thereby subverting this tightly controlled balance. A central mechanism enabling this transformation is the dysregulation of growth factor receptors, exemplified by the overexpression of EGFR in many cancers, which renders cancer cells hyperresponsive to ambient concentrations of growth factors that would otherwise be inadequate to induce proliferation^40,41^. More importantly, TMEM132A circumvents normal regulatory constraints by persistently redirecting internalized EGFR to recycling endosomes and back to the plasma membrane, thereby bypassing lysosomal degradation and "locking" the EGFR signaling pathway into a constitutively active state-a process that operates independently of EGF stimulation. This mechanistic rerouting fundamentally alters the dynamics of EGFR signaling, driving sustained plateau-phase activation of PI3K/AKT cascades, including lipid synthesis pathway, rather than the transient pulse-like responses characteristic of ligand-induced signaling. Such sustained activation not only confers a proliferative advantage under growth factor-limiting conditions but also diminishes the efficacy of therapeutic agents that rely on receptor internalization and degradation, while the EGF-independent nature of this process enables TMEM132A to operate autonomously within the tumor microenvironment, contributing to intratumoral heterogeneity and resistance to conventional EGFR-targeted therapies. Collectively, these observations position TMEM132A as a master regulator that reconfigures EGFR circuitry, suggesting that targeting TMEM132A or the recycling endosome machinery it exploits may offer a more effective strategy for dismantling the growth signal autonomy that drives tumor progression across diverse cancer types.

EGFR T790M mutation is the principal mechanism underlying acquired resistance to first-and second-generation EGFR tyrosine kinase inhibitors (EGFR-TKIs) in NSCLC. Although third-generation EGFR-TKIs such as osimertinib effectively target T790M, post-treatment resistance remains inevitable, often characterized by either the loss of T790M or the emergence of additional mutations such as C797S^42^. This evolving resistance landscape suggests that targeting individual kinase domain mutations is insufficient to counteract the adaptive and evasive capabilities of tumor cells. Therefore, a paradigm shift from "mutation-specific inhibition" to "global suppression of EGFR signaling" is warranted. LFNanoT132A#3 offer distinctive mechanistic advantages in overcoming T790M-mediated resistance. First, LFNanoT132A#3 enable reprogramming of protein homeostasis: TMEM132A-targeting nanobodies disrupt the aberrant TMEM132A–EGFR interaction, redirect EGFR trafficking from recycling to lysosomal degradation, thereby achieving global downregulation of EGFR protein levels. This mechanism operates independently of the mutational status of the kinase domain, rendering it theoretically effective against both T790M single mutations and T790M/C797S compound mutations, and fundamentally precludes point mutation-driven resistance escape. Second, nanobodies exert their function through binding to extracellular domains, a mode of action independent of ATP-binding pocket conformation; hence, they are unaffected by steric changes induced by T790M or C797S mutations, bypassing the core limitation of TKIs whose efficacy is compromised by structural alterations in the target. This degradation-based, global downregulation strategy represents a departure from the conventional "mutation chase" with cancer cells. As such, LFNanoT132A#3 offers a potentially more durable therapeutic approach for patients with T790M-associated resistance.

However, several mechanistic aspects of our study remain to be explored. KEGG pathway analysis of transcriptomic data revealed that TMEM132A regulates classical downstream effectors of EGFR signaling, including the PI3K/AKT, mTOR, and MAPK pathways, with the most pronounced effect observed on the PI3K pathway, whereas no significant regulation of the JAK/STAT or PLCγ/PKC pathways was detected (Fig. 5A). The mechanistic basis underlying the selectively interfering of PI3K/AKT by TMEM132A presents an interesting and worthwhile question for further investigation. Additionally, we found that TMEM132A also modulates the transmembrane receptor protein serine/threonine kinase signaling pathway as well as the WNT pathway (Fig. 3A). Although a direct regulatory role of TMEM132A on the Wnt pathway has been reported in 293T cells, our IP-MS results did not detect any interaction between TMEM132A and Wnt pathway-related proteins. Therefore, how TMEM132A regulates the Wnt pathway in the context of tumor progression warrants further exploration.

## Materials & Methods

### Cell culture

786-O, RCC10, RCC4, UMRC-2, A498, UMRC6 BT-474, MCF-7, MDA-MB-231, T98G, Huh7, A875, LNCaP and A549 cells were cultured in DMEM (Gibco, 11965118) supplemented with 10% FBS and 1% penicillin–streptomycin. 769-P, BxPC3, H1975 and HCC827 cells were cultured in RPMI-1640 (Gibco, 11875093) with 10% FBS and 1% penicillin–streptomycin. SK-N-BE2 cells were cultured in DMEM-F12 (Gibco, 11320033) supplemented with 10% FBS and 1% penicillin–streptomycin. Mycoplasma contamination was routinely monitored by PCR and treated with Myco-Zero™ Plus (Beyotime, C0280).

### Plasmid construction and Cell transfection

For stable overexpressing proteins, the open reading frame (ORF) sequences of TMEM132A and EGFR were cloned into the phage-CMV-3×FLAG vector with FLAG tag at the C-terminus. To transient express proteins, full length and truncated fragments (amino acids 392-635) of TMEM132A were clone into the pcDNA3.1 vector with an HA-tag at the N-terminus. The truncated EGFR fragment (25-645 aa) was inserted into the pcDNA3.1 vector with his-tag at the C-terminus. Mutant TMEM132A plasmids were generated based on phage-FLAG-TMEM132A plasmid using site-directed mutagenesis kit (Vazyme, Cat: C216-01). For knock down of TMEM132A in cancer cells, two independent shRNAs were cloned into plKO.1-puro or Tet-on plKO.1-puro plasmid. For TMEM132A CRISPR knockout in cancer cells, two distinct sgRNAs sequences were inserted into the lentiGuide-Puro plasmid (Addgene #52963). The sequences of primers for plasmid construction are listed in Table 1.

**Table 1.**
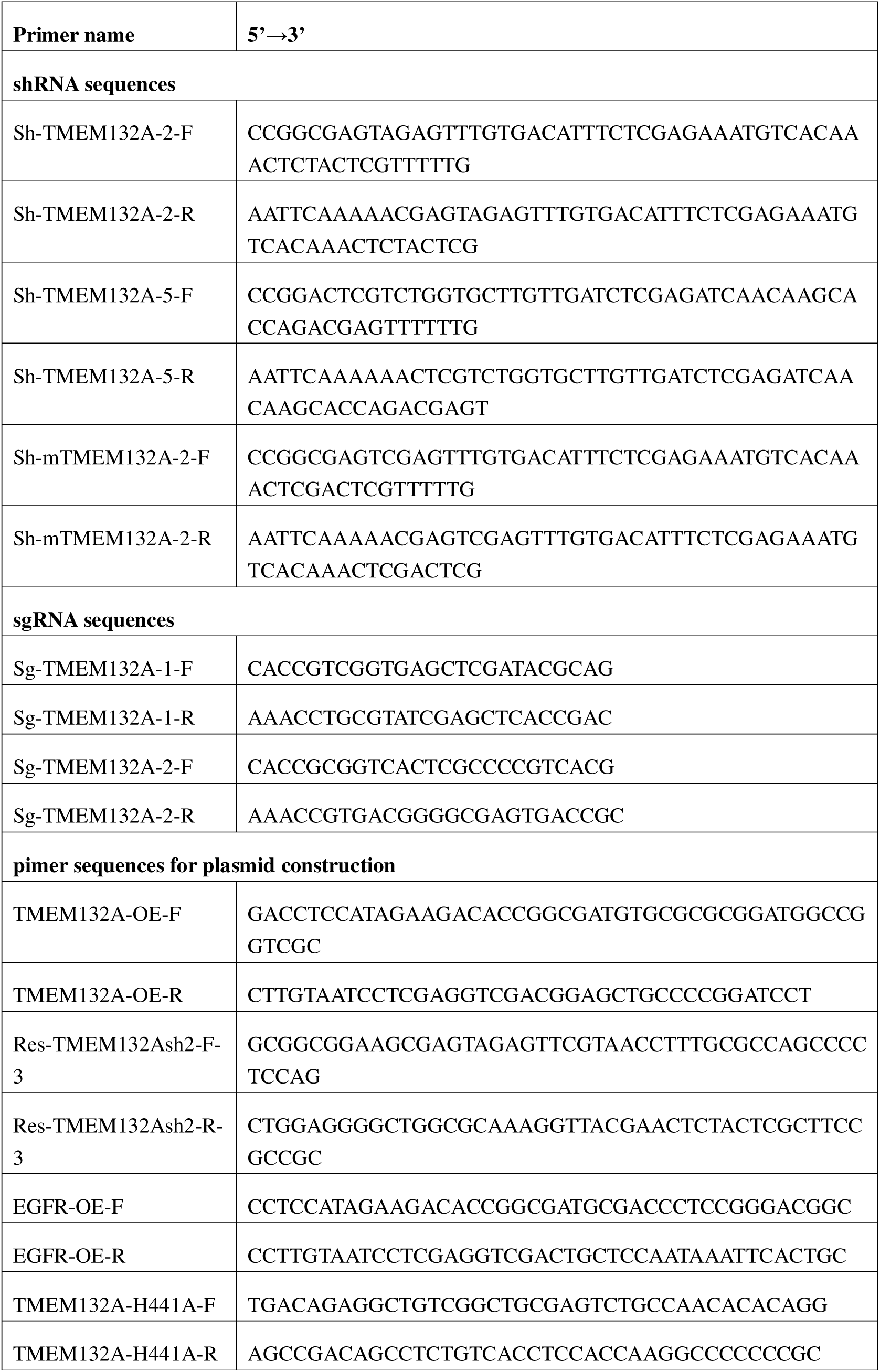
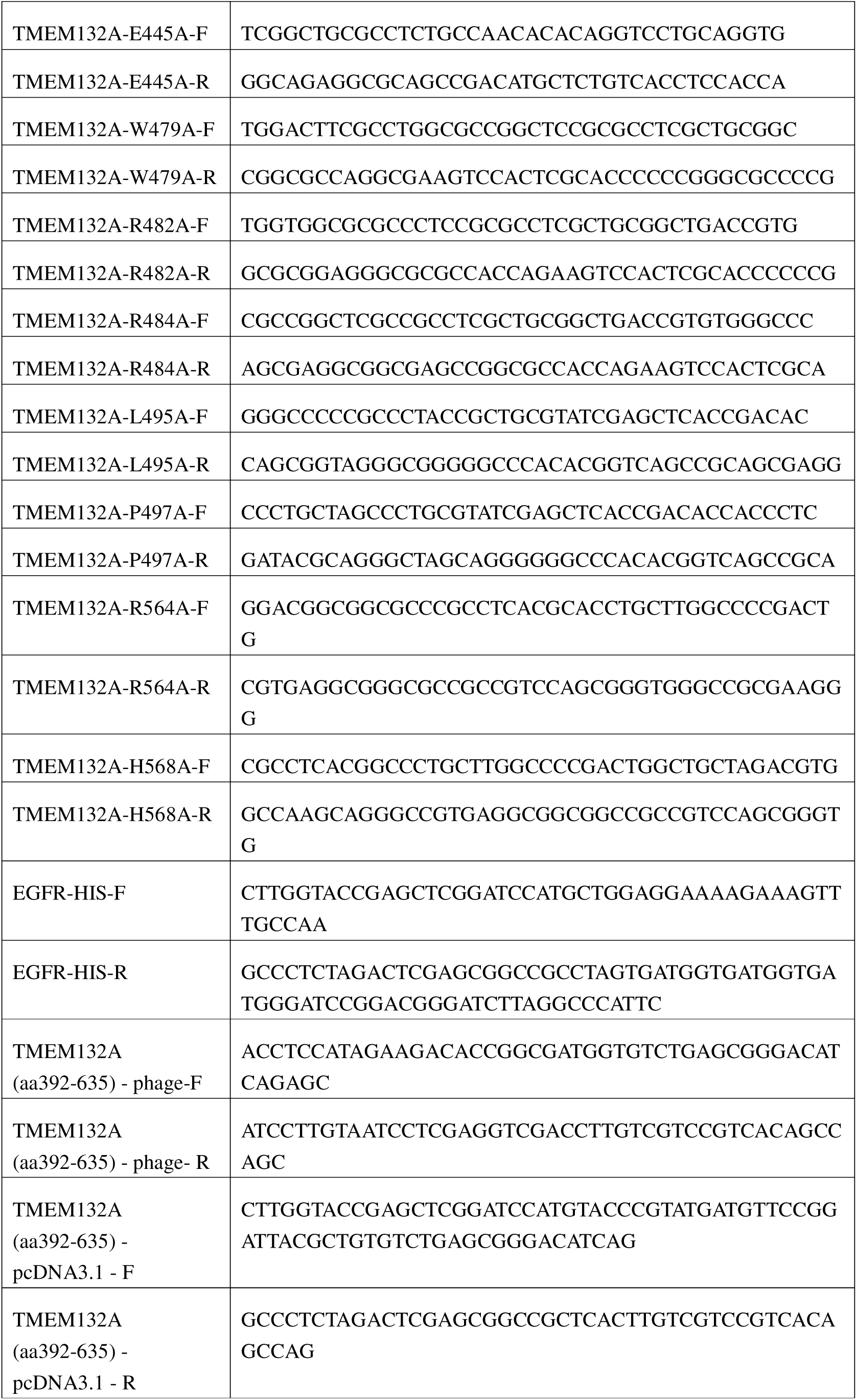
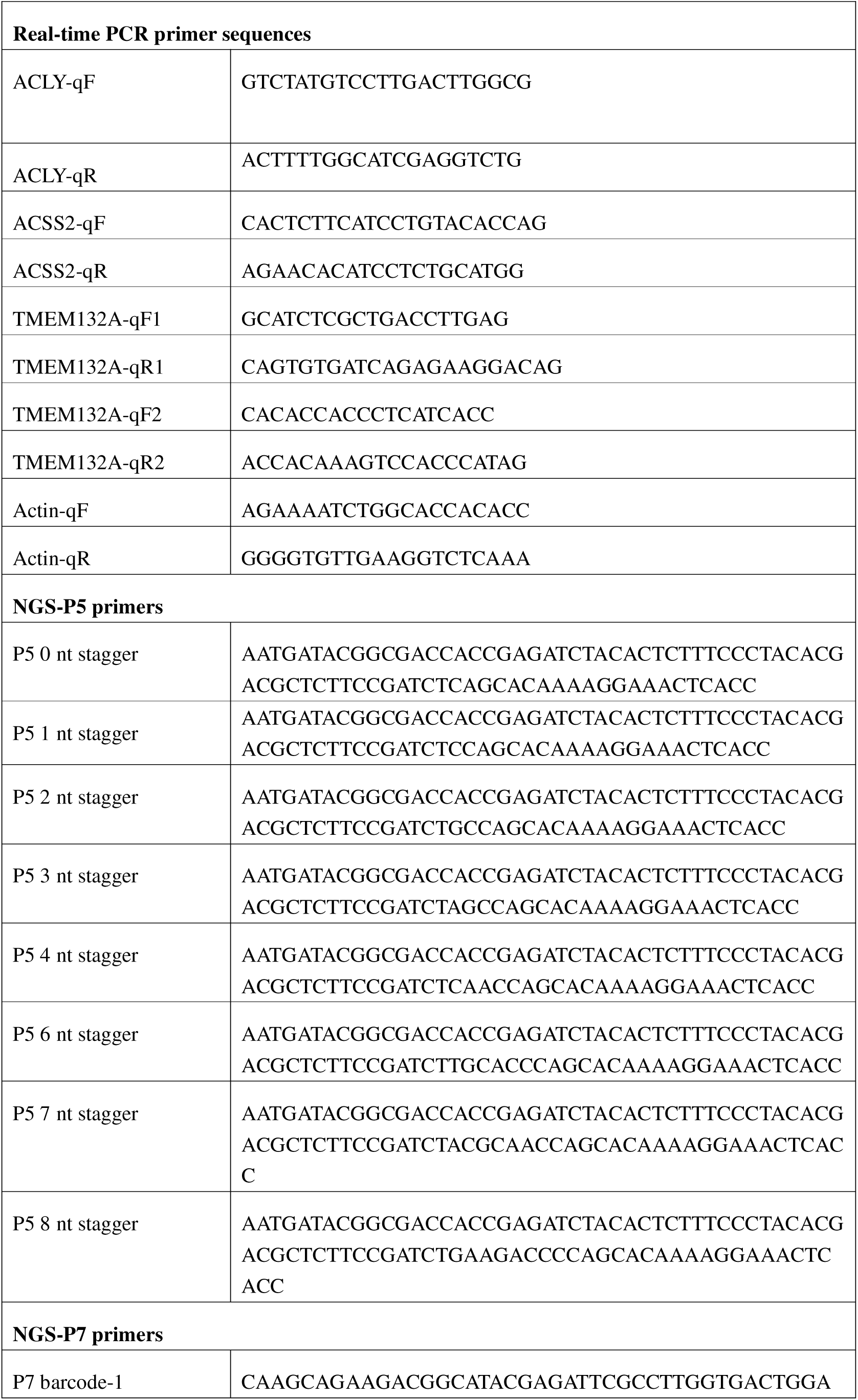
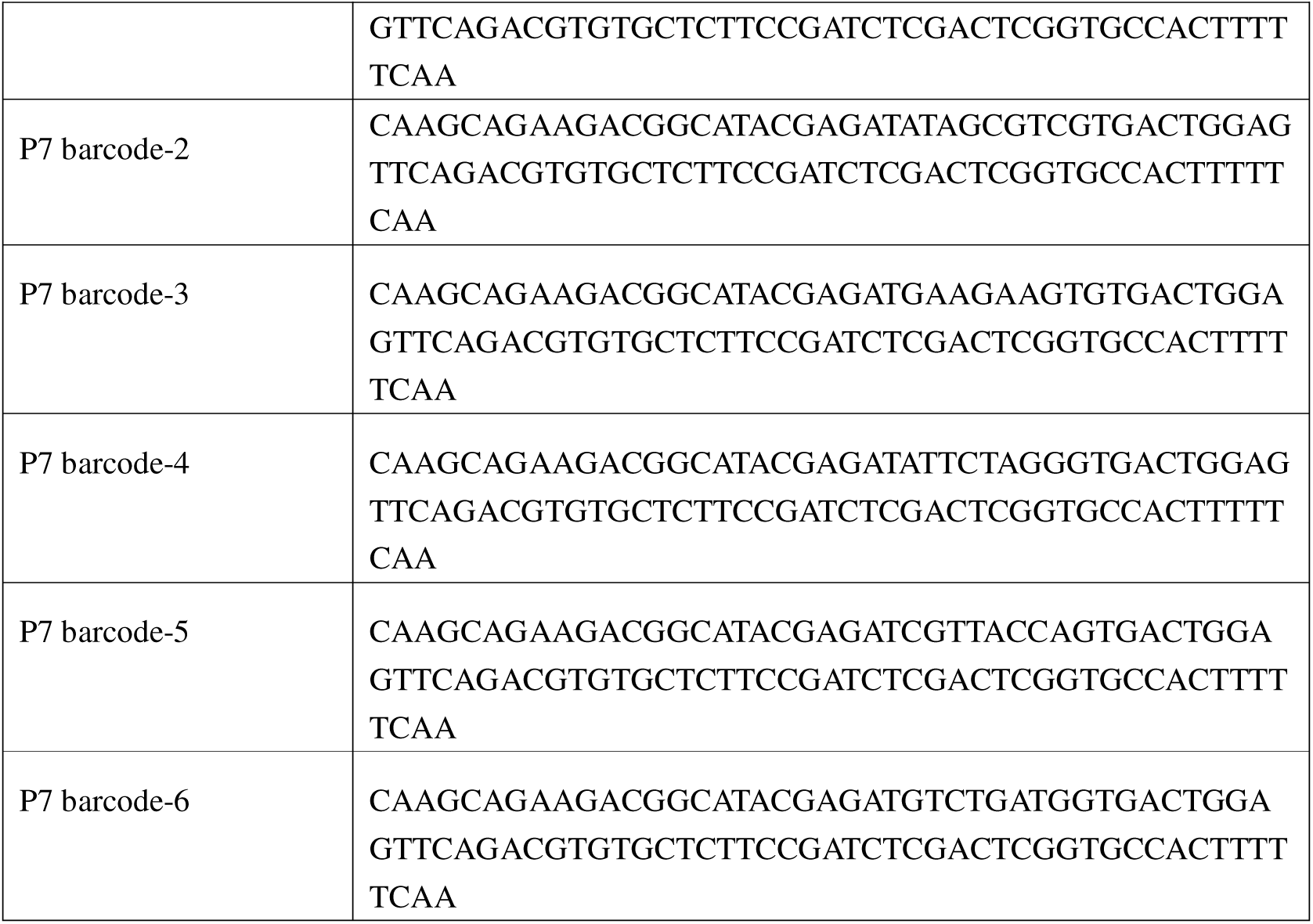
shRNA sequences, sgRNA sequences, real-time PCR primer sequences and pimer sequences for plasmid construction and Crispr cas9 screening.

Transient transfections of 293T cells were conducted using poluethylenimine (PEI) MW 4000 (Yeasen, 40816ES01) following the manufacture’s instruction. The plasmids-PEI mixture at a weight ratio of 3:1 was added to the cell medium. For lentiviruses packaging, lentiviruses were prepared by transfecting 293T cells as previously described^23^.

### Stable cell lines generation

Different cancer cells were transduced by lentiviruses to generate stable overexpression or knock down/out cells. Cells were seeded in 6 well plate at a density of 1.5×10^5^ cells per well. Lentivirus were added into cells with polybrene (8μg/mL). Transduced cells were selected at a desired antibiotic concentration, refresh the selection antibiotic every 2 days. For the Tet-on inducible knockdown of TMEM132A, stable cells were incubated with 2μg/ml doxycycline for at least 72 h. The efficiency of overexpression and knockdown/ out was confirmed by immunoblotting.

### CRISPR screening and analysis

A human CRISPR deletion library - trafficking, mitochondrial (addgene#101931) was re-amplified using liquid culture method according to manufacturer’s instruction and the titer of sgRNA library was determined in 293T cells as previous reproted^23^. A total of 4.0×10^7^ 786-O cells with stable Cas9 expression were infected with the sgRNA library at a multiplicity of infection (MOI) <0.3. Three days after infection, cells were subjected to puromycin (1μg/mL) selection for an additional 5 days and then harvested to serve as the baseline sample. The remaining selected cells were cultured for additional 14 days, with the medium refreshed every 3 days, and then harvested at the endpoint. Genomic DNA (gDNA) was extracted using the ZYMO genomic DNA extraction kit (D4075). The sgRNA-encoding sequences were amplified by PCR using high-fidelity Taq polymerase (Yeasen, 10153ES). The sequences of primers for sgRNA amplification are listed in Table 1.

The computational analysis of CRISPR-Cas9 screening data was performed using MAGeCK (version 0.5.9.5)^43,44^. Raw sequencing reads in FASTQ format were processed using the “mageck count” module to quantify sgRNA abundance by mapping sequencing reads to the sgRNA reference sequences of the Human CRISPR Deletion Library-Trafficking, mitochondrial (Addgene #101931). To identify negatively selected genes whose sgRNAs were significantly depleted during the screen, the “mageck test” module was applied. Read counts were normalized using non-targeting control sgRNAs to correct for differences in sequencing depth and potential library composition bias. Gene-level depletion analysis was performed using the Robust Rank Aggregation (RRA) algorithm, which integrates signals from multiple sgRNAs targeting the same gene. Statistical significance was determined by calculating the false discovery rate (FDR), and genes with FDR < 0.05 were considered significant hits.

### Cell Proliferation and Anchorage-Independent Growth assay

Cells were seeded into 96-well plates at a density of 1000 cells per well, and cell viability was analyzed using CCK-8 at indicated time points as the manufacture’s protocol (Beyotime, C0040). For colony formation ability, cells were seeded into 6-well plates at a density of 2000 cells per well. When cell colonies in the control group reached a suitable density and then were fixed with 4% paraformaldehyde and stained with crystal violet. For the anchorage-independent growth assay (3-D soft agar assay), 786-O cells were cultured at a density of 40,000 cells/mL, while other cells were cultured at 30,000 cells/mL as described previously^45^.

### Immunoblots and co-immunoprecipitation (Co-IP)

For immunoblots assay, cell pellets were lysed in EBC buffer (0.1mM EDTA, 50mM Tris-HCl, 120mM NaCl, 0.5% NP-40 and 10% Glycerol) and tissues were lysed in Urea (8mM) supplemented with protease (Roche, 04693116001) and phosphatase inhibitor (Roche, 04906837001). After sonication and centrifugation, supernatants were collected, subjected to SDS-PAGE and immunoblotted with indicated antibody. Proteins were transferred onto Nitrocellulose membrane (Millipore).

Co-IP was performed to confirm endogenous and exogenous protein-protein interactions. Briefly, primary antibody was added into cell lysate and incubated at 4°C overnight with gentle rotation. Lysate incubated with antibodies was added to 10 μl of protein G agarose beads (Roche Applied Bioscience, 11243233001) with rotation for an additional 3 hours at 4°C. The bound complexes were then centrifuged, washed with EBC buffer, subjected to SDS-PAGE, and immunoblotted with indicated antibody. Antibodies and the dilution ratios used in this work were list in Table 2.

**Table 2.**
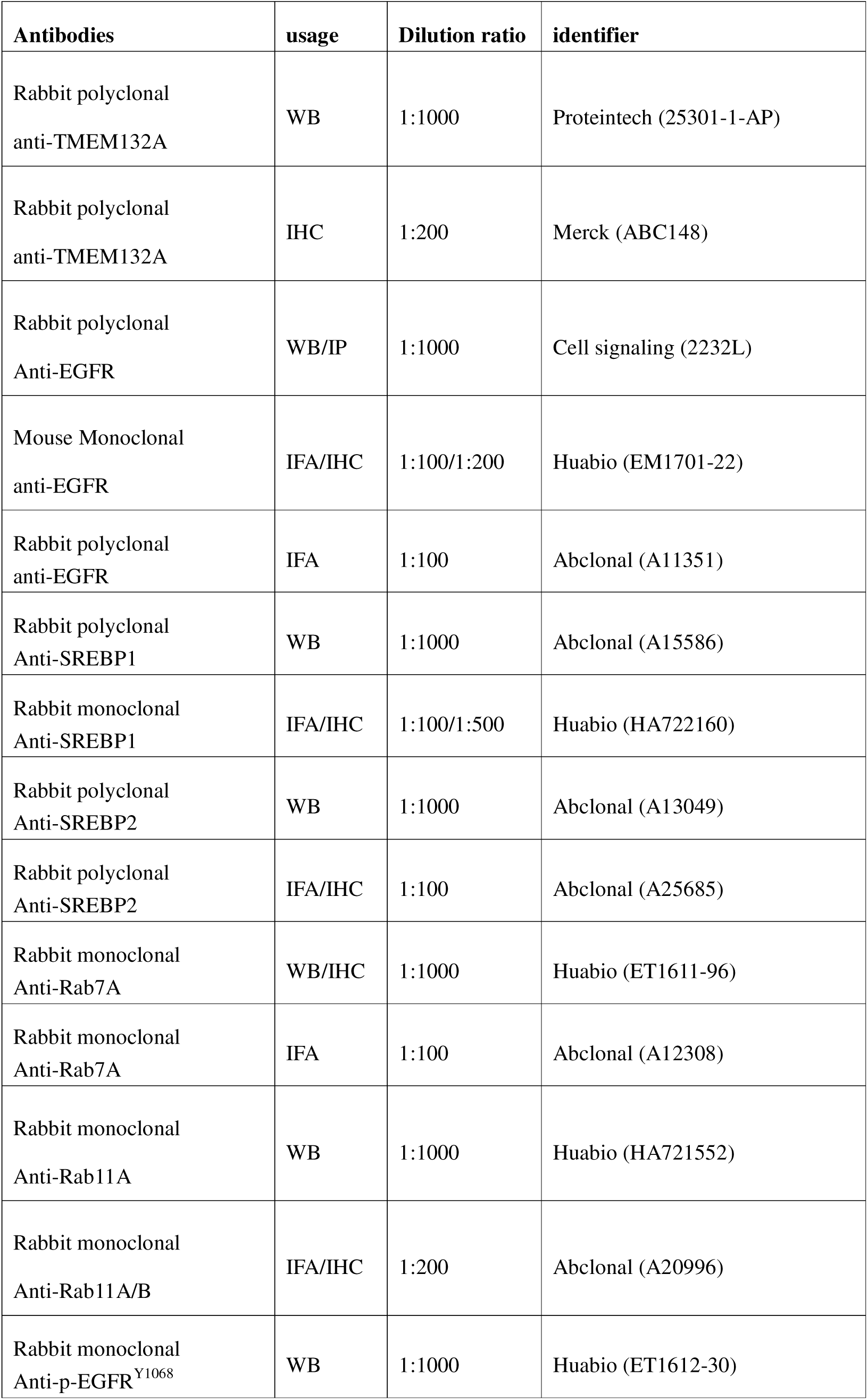
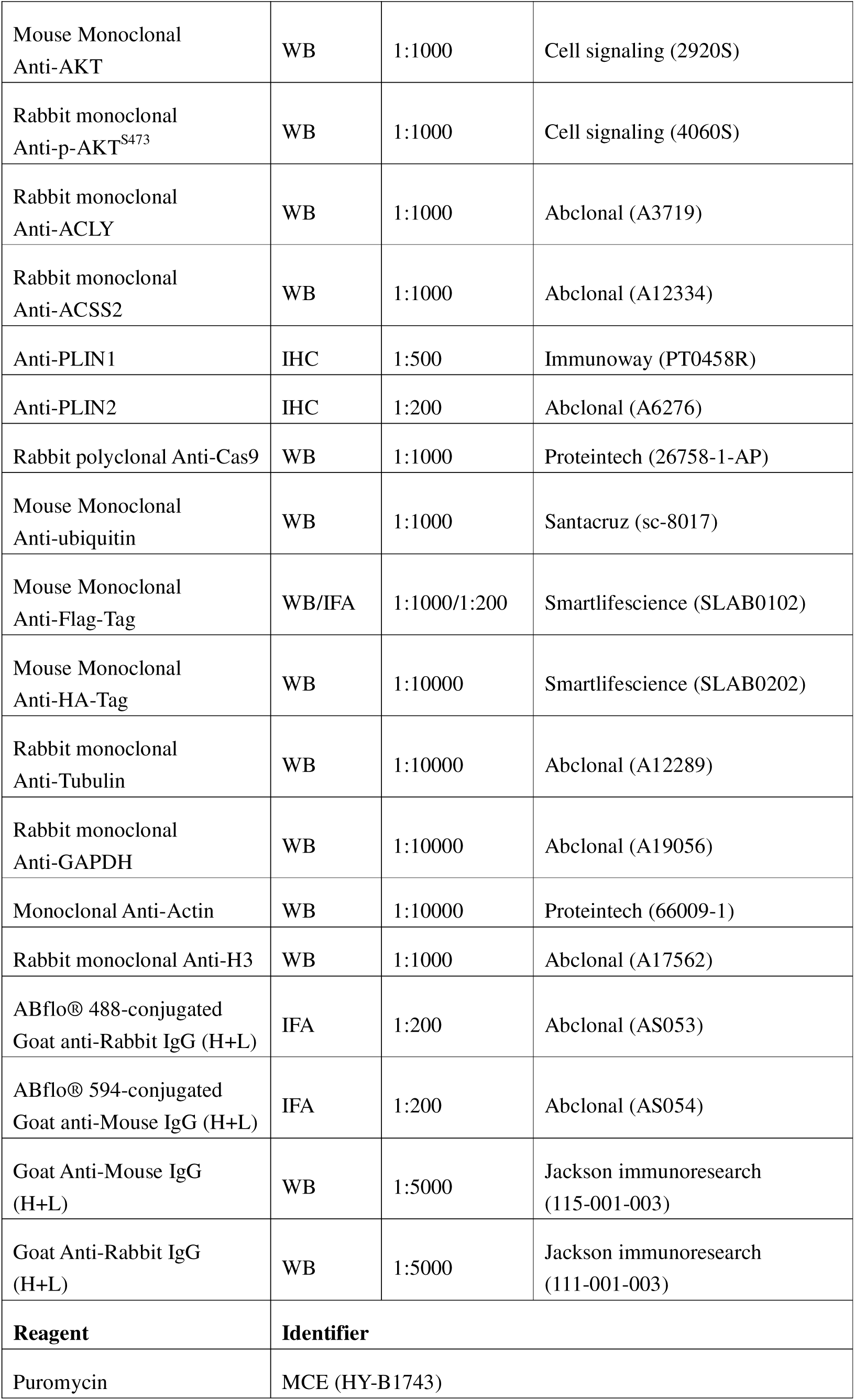
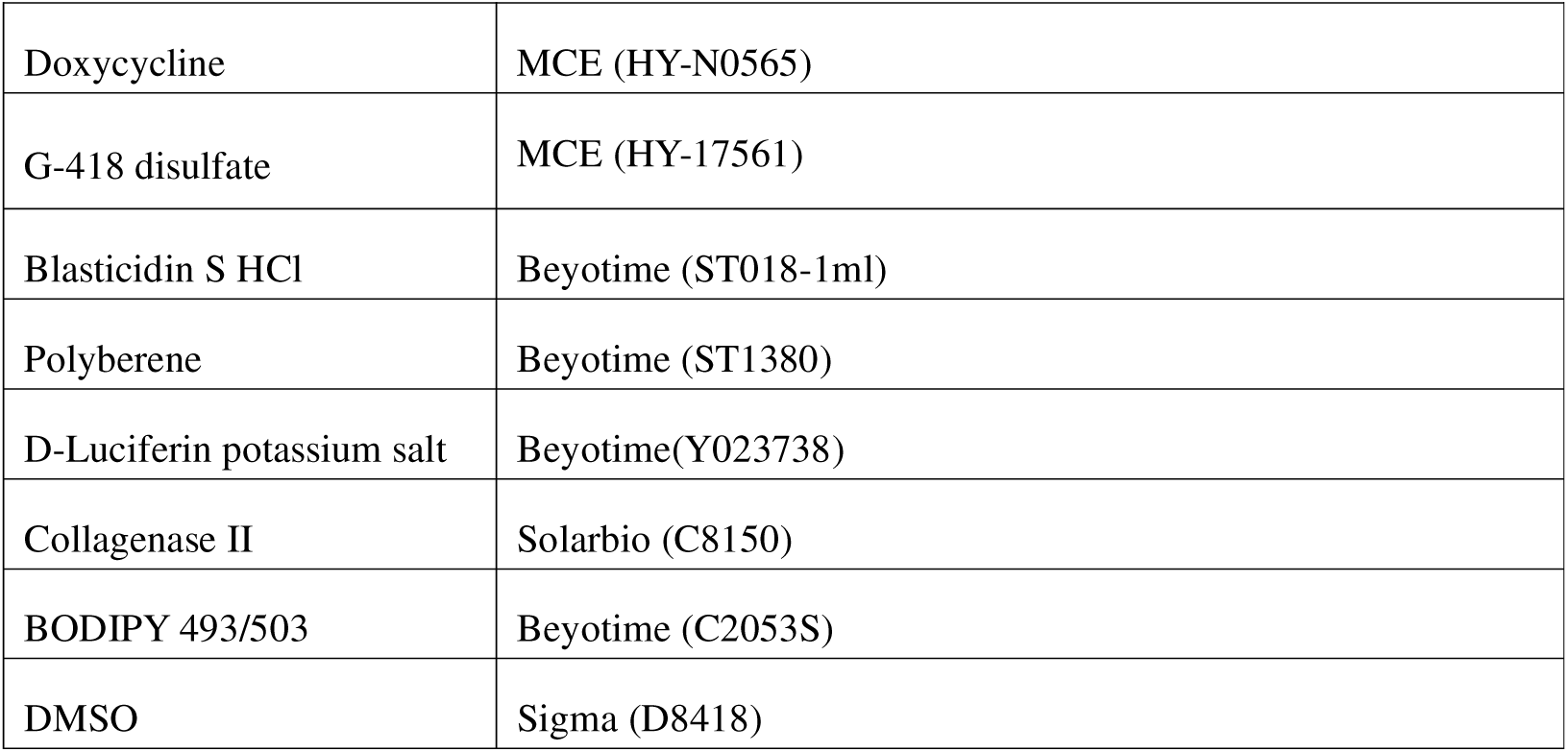
Summary of antibodies.

| **Table 2 Summary of antibodies.** |  |  |  |
| --- | --- | --- | --- |
| **Antibodies** | **usage** | **Dilution ratio** | **identifier** |
| Rabbit polyclonal<br>anti-TMEM132A | WB | 1:1000 | Proteintech (25301-1-AP) |
| Rabbit polyclonal<br>anti-TMEM132A | IHC | 1:200 | Merck (ABC148) |
| Rabbit polyclonal<br>Anti-EGFR | WB/IP | 1:1000 | Cell signaling (2232L) |
| Mouse Monoclonal<br>anti-EGFR | IFA/IHC | 1:100/1:200 | Huabio (EM1701-22) |
| Rabbit polyclonal<br>anti-EGFR | IFA | 1:100 | Abclonal (A11351) |
| Rabbit polyclonal<br>Anti-SREBP1 | WB | 1:1000 | Abclonal (A15586) |
| Rabbit monoclonal<br>Anti-SREBP1 | IFA/IHC | 1:100/1:500 | Huabio (HA722160) |
| Rabbit polyclonal<br>Anti-SREBP2 | WB | 1:1000 | Abclonal (A13049) |
| Rabbit polyclonal<br>Anti-SREBP2 | IFA/IHC | 1:100 | Abclonal (A25685) |
| Rabbit monoclonal<br>Anti-Rab7A | WB/IHC | 1:1000 | Huabio (ET1611-96) |
| Rabbit monoclonal<br>Anti-Rab7A | IFA | 1:100 | Abclonal (A12308) |
| Rabbit monoclonal<br>Anti-Rab11A | WB | 1:1000 | Huabio (HA721552) |
| Rabbit monoclonal<br>Anti-Rab11A/B | IFA/IHC | 1:200 | Abclonal (A20996) |
| Rabbit monoclonal<br>Anti-p-EGFR<sup>Y1068</sup> | WB | 1:1000 | Huabio (ET1612-30) |
| Mouse Monoclonal<br>Anti-AKT | WB | 1:1000 | Cell signaling (2920S) |
| Rabbit monoclonal<br>Anti-p-AKT<sup>S473</sup> | WB | 1:1000 | Cell signaling (4060S) |
| Rabbit monoclonal<br>Anti-ACLY | WB | 1:1000 | Abclonal (A3719) |
| Rabbit monoclonal<br>Anti-ACSS2 | WB | 1:1000 | Abclonal (A12334) |
| Anti-PLIN1 | IHC | 1:500 | Immunoway (PT0458R) |
| Anti-PLIN2 | IHC | 1:200 | Abclonal (A6276) |
| Rabbit polyclonal Anti-Cas9 | WB | 1:1000 | Proteintech (26758-1-AP) |
| Mouse Monoclonal<br>Anti-ubiquitin | WB | 1:1000 | Santacruz (sc-8017) |
| Mouse Monoclonal<br>Anti-Flag-Tag | WB/IFA | 1:1000/1:200 | Smartlifescience (SLAB0102) |
| Mouse Monoclonal<br>Anti-HA-Tag | WB | 1:10000 | Smartlifescience (SLAB0202) |
| Rabbit monoclonal<br>Anti-Tubulin | WB | 1:10000 | Abclonal (A12289) |
| Rabbit monoclonal<br>Anti-GAPDH | WB | 1:10000 | Abclonal (A19056) |
| Monoclonal Anti-Actin | WB | 1:10000 | Proteintech (66009-1) |
| Rabbit monoclonal Anti-H3 | WB | 1:1000 | Abclonal (A17562) |
| ABflo® 488-conjugated<br>Goat anti-Rabbit IgG (H+L) | IFA | 1:200 | Abclonal (AS053) |
| ABflo® 594-conjugated<br>Goat anti-Mouse IgG (H+L) | IFA | 1:200 | Abclonal (AS054) |
| Goat Anti-Mouse IgG<br>(H+L) | WB | 1:5000 | Jackson immunoresearch<br>(115-001-003) |
| Goat Anti-Rabbit IgG<br>(H+L) | WB | 1:5000 | Jackson immunoresearch<br>(111-001-003) |
| **Reagent** | **Identifier** |  |  |
| Puromycin | MCE (HY-B1743) |  |  |

|  |  |
| --- | --- |
| Doxycycline | MCE (HY-N0565) |
| G-418 disulfate | MCE (HY-17561) |
| Blasticidin S HCl | Beyotime (ST018-1ml) |
| Polyberene | Beyotime (ST1380) |
| D-Luciferin potassium salt | Beyotime(Y023738) |
| Collagenase II | Solarbio (C8150) |
| BODIPY 493/503 | Beyotime (C2053S) |
| DMSO | Sigma (D8418) |

To assess the blocking effect of nanobodies LFNanoT132A#1 and #3 on the TMEM132A-EGFR interaction, cell lysates were pre-incubated with control nanobody (NanoIgG) or LFNanoT132A#1 or #3 overnight at 4°C with gentle agitation. On the following day, the EGFR antibody was introduced into the lysates, and the IP was subsequently performed according to the standard protocol.

### LC-MS/MS analysis

Cell lysates from 786-O control (EV) cells and 786-O FLAG-TMEM132A (OE) cells were subjected to immunoprecipitation (IP) using anti-FLAG M2 affinity agrose (Sigma-Aldrich, A2220). Immunoprecipitates were separated by SDS-PAGE and visualized by Coomassie staining. Then the gel strips of EV and OE group were pretreated and mass spectrometry analysis by Shanghai Bioprofile Co., LTD. For MS analysis, an appropriate number of peptides from each sample was subjected to chromatographic separation using the nano-flow Easy nLC 1200 Chromatography System. After peptide separation, data-dependent acquisition (DDA) MS analysis was performed on the Q-Exactive HF-X Mass Spectrometer (Thermo Scientific). Full MS resolutions were set to 60,000 at m/z 200 and mass range was set to 350–1800 m/z. MaxQuant (version 2.4.14.) software was utilized for database searching and protein identification.

### Ubiquitination analysis

Cells were harvested and lysed in 100ul EBC buffer containing 1% SDS. Cell extraction were heat-denatured for 5 min at 95°C to disrupt non-covalent protein-protein interactions and then diluted with 900μL EBC buffer. After sonication and centrifugation, a small portion of supernatant was collected as an input sample and the majority of cell lysates were immunoprecipitated with EGFR antibody (Cell Signaling Technology, 2232L). The immunoprecipitated proteins were subjected to SDS-PAGE, and then analyzed using immunoblotting with ubiquitin antibody (Santa Cruz Biotechnology, sc-8017).

### Immunofluorescence

Immunofluorescence assays were performed as previously reported^46^. Briefly, cells were seeded onto coverslips placed in 12 well plate at a density of 5×10^4^ cells per well and cultured overnight to allow adherence. Cells were fixed using 4% paraformaldehyde for 15 min, permeabilized with 1% Triton X-100 for 15min at room temperature and then blocked by 30% FBS at least 10 min. Cells were then incubated with corresponding primary antibody at 4°C overnight and fluoresent-conjuncted secondary antibody at room temperature for 1 h in dark. Coverslips were mounted with an antifade solution containing 4,6-diamidino-2-phenylindole (DAPI). Images were captured using a laser confocal fluorescence microscopy instrument (Zeiss LSM 900) with a 60 × oil immersion objective.

### Acetyl-CoA Measurement

Acetyl-CoA in cells and tissues were determine using Acetyl-CoA assay kit (Solarbio, BC0980) as the manufacture’s instruction. Briefly, 3×10^6^ cells were harvested and lysed in 200μL extraction buffer on ice for 30 min, followed by sonication for 4s at 80 watts 50 times with 5s interval. For tissues, 0.05g tumor samples were scissored into small pieces and immerged in 500 μL extraction buffer and homogenized on ice thoroughly. After centrifugation at 11,000 × g for 10 min at 4°C, the supernatants were collected for quantification of total Acetyl-coA levels. The absorbance was detected with a spectrophotometer at 340nm.

### Metabolic profiling

For metabolic profiling, metabolites from 786-O Ctrl cells (shCtrl) and TMEM132A-depleted cells (TMEM132A-sh5) were extracted using pre-cooled mixtures of methanol, acetonitrile and water (v/v/v, 2:2:1). Samples were ultrasonic shaked on ice baths 1 h and centrifuged at 14000×g for 20 min at 4°C. The supernatant was collected and concentrated to dryness in vacuum. Dried metabolites were dissolved in 50% acetonitrile and then analyzed using the UHPLC-ESI-Q-orbitrap-MS system (UHPLC, Shimadzu Nexera X2 LC-30AD, Shimadzu, Japan) coupled with Q-Exactive Plus (Thermo Scientific, San Jose, USA). Multivariate statistical analysis of metabolomic data was performed using partial least squares discriminant analysis (PLS-DA) and orthogonal partial least squares discriminant analysis (OPLS-DA) to achieve data dimensionality reduction, group discrimination, and removal of noise. Variable importance in the projection (VIP) scores were calculated based on the OPLS-DA model with a threshold of VIP > 1.0 indicating potential biomarkers. For two-group comparisons, Student’s t-test and fold change (FC) analysis were used. Metabolites with VIP > 1 and P < 0.05 were defined as significantly differential metabolites. If OPLS-DA modeling was unsuccessful, thresholds of [FC] ≥ 1.5 combined with P < 0.05 were applied. Differential metabolites were classified according to their chemical structures and functions. Functional categorization of metabolites was performed using HMDB database (Super Class and Class), as well as Ontology annotations obtained from the ClassyFire website (http://classyfire.wishartlab.com).

### RNA isolation, qRT-PCR and RNA-seq

Total cellular RNAs were extracted using Quick-DNA/RNA Microprep Plus Kit (Zymo research, R1055). RNA (1μg) was reverse-transcribed using the ABScript III RT Master Mix for qPCR with gDNA Remover (Abclonal, RK20429). QRT-PCR was performed on an ABI QuantStudio 5 real-time PCR system. Relative mRNA expression level was calculated by the 2-ΔΔCt method and normalized to the endogenous gene Actin. Primers used in qRT-PCR are listed in Table 1.

Total RNA (500ng) was used to construct libraries using the Hieff NGS® mRNA Isolation Master Kit V2 mRNA for Illumina (Yeasen, 12629ES) following the manufacturer’s instructions. Sequencing was performed on a NovaSeq 6000 platform (Illumina) by HaploX Biotechnology Co., LTD. The raw sequencing data were preprocessed using fastp (v0.23.4)^47^ to remove adapter sequences and filter out low-quality reads. The resulting high-quality clean reads were aligned to the human reference genome (GRCh38) using HISAT2 (v2.2.1)^48^. Alignment files were then sorted and converted to BAM format via samtools (v1.9). To quantify gene expression, the number of reads mapped to each gene was calculated using featureCounts (v2.0.6)^49^ based on the Ensembl human genome annotation (Homo_sapiens.GRCh38.90.gtf). Differential expression analysis was performed using the DESeq2 (v1.42.1)^50^ package in R (v4.3.3). To further investigate the biological significance of the identified differentially expressed genes (DEGs), Gene Ontology Biological Process (GOBP) enrichment analysis was conducted using the clusterProfiler (v4.10.1)^51^ package. A p-value cutoff of 0.05 was applied to identify significantly enriched biological pathways.

### Patient-derived cancer organoids (PDOs)

The construction of breast cancer organoids was performed following previously described protocols^52,53^. For knockdown of TMEM132A in the organoids, organoids were dissociated and incubated with high-titer infectious shTMEM132A lentivirus at an MOI of 5 at 37°C for 30–60 min. The organoids were then plated on Matrigel (Corning, 356255) and cultured in human complete feeding medium (Mogengel, MA-0807T011LP). Three days after transduction, expansion medium containing 1μg/mL puromycin was used for organoid culture, with the medium refreshed every 3 days.

### Patient-derived Xenograft (PDX)

Fresh tissues were cut into 3–5 mm fragments and implanted into the subrenal capsule pocket of the kidneys of eight-week-old NSG mice. Tumor growth was monitored by palpation every week. Approximately 3 to 4 months after surgery, the tumor tissues were passaged into subsequent cohorts once the tumor diameter reached at least 7 mm. The passaged tumor tissues were analyzed by hematoxylin and eosin (H&E) staining to confirm the cancer type.

### PDX cell line isolation

Tumor tissues from PDX mice were minced into small pieces using a scalpel blade and then incubated in digestion mixture (DMEM supplemented with 1% penicillin–streptomycin, 5% FBS, and collagenase I) in a shaker at 37°C for 2 h. The dispersed cells were filtered through a 40μm cell strainer and treated with red blood cell lysis buffer (Sigma, 11814389001). The remaining cells were resuspended and cultured in complete DMEM medium. Cells were passaged for at least 10 generations to obtain a stable PDX-derived cell line, which was then used for functional investigation of TMEM132A.

### Orthotopic tumor Growth

For ccRCC, approximately 2×10^6^ 786-O cells were resuspended in 20 µL fresh medium and 20 µL Matrigel (Corning, 354248) and injected orthotopically into the left kidney of eight-week-old NOD/ShiLtJGpt-*Prkdc^em26Cd52^Il2rg^em26Cd^*^22^/Gpt (NCG) mice. Bioluminescence imaging was performed to confirm successful tumor engraftment in the kidney. Mice were given drinking water containing doxycycline (2 mg/L). Mice were euthanized 5 weeks after doxycycline treatment.

For TNBC, approximately 2×10^6^ MDA-MB-231 cells or 1×10^6^ PDX-derived tumor cells were resuspended in 50 µL fresh medium and 50 µL Matrigel and injected orthotopically into the mammary fat pads of female NCG mice. Tumor volume was measured with digital calipers every 7 days for approximately 6 weeks (for MDA-MB-231 cells) after tumors became palpable, or at week 8 after transplantation of PDX-derived TNBC tumor cells. Tumor volume was calculated using the formula: 0.5 × L × W², where L is the larger diameter and W is the smaller diameter.

### Cell line derived xenograft models

For NSCLC allografts, about 5×10^6^ A549 or H1975 cells were resuspended in 50 μL fresh medium and 50 μL matrigel and subcutaneously injected into the right flank of NCG mice.

### Tumor tissue derived syngeneic models

For breast cancer, fresh tumor tissues from 4T1 murine mammary carcinoma cell-bearing mice were cut into 3–5 mm fragments and orthotopically implanted into the mammary fat pad of female BALB/c mice (8 weeks old). For pancreatic cancer, tumor tissues from Panc02 cell-bearing mice were implanted subcutaneously into C57BL/6J mice (8 weeks old) using the same procedure.

### Drug treatment

For EGF treatment, cells were serum-starved for 5 h and then treated with 10 ng/mL EGF (MCE, HY-P7109). For other inhibitor treatments, cells were treated with CHX (MCE, HY-12320), chloroquine (MCE, HY-17589A), MG132 (MCE, HY-13259), Ac-CoA Synthase Inhibitor 1 (MCE, HY-104032), bempedoic acid (MCE, HY-12357), or sodium dichloroacetate (MCE, HY-Y0445A) at the indicated concentrations as labeled in the figures or figure legends.

### Cell Fractionation

Cell fractionation were performed as previously reported^54^. Cell pellet was washed with ice-cold PBS, resuspended in cell lysis buffer (buffer A, 10Mm HEPES, [pH, 7.9], 10mM KCl, 1.5mM MgCl_2_, 0.34 M sucrose, 10% glycerol, 1m M DTT and 0.1% Triton X-100) with 1% protease and phosphatase inhibitors and lysed for 15 min on ice. The suspensions were collected to a new vial as whole cell lysate (WCL). Then the solution were centrifuged at 1500×g for 10 min at 4°C. Pellet was collected and washed six times in buffer A. Nuclei were collected by high-speed centrifugation (12000×g, 5 min, 4°C) and lysed in buffer B (3mM EDTA, 0.2mM EGTA, 1m M DTT) with 1% protease and phosphatase inhibitors.

### Immunohistochemistry (IHC) and multiplexed IHC (mIHC)

IHC staining was performed using the SABC (Rabbit) IHC Kit (Biosharp) according to the manufacturer’s instructions. Paraffin-embedded mouse tumor tissues or patient samples were sectioned into 5-μm-thick slices, incubated in citrate buffer (pH 6.0) for 5 min at 120°C, and treated with 0.3% H_2_O_2_ for 10 min to block endogenous peroxidase activity. The sections were then incubated with normal goat serum (Zsbio, ZLI-9056) for 10 min at room temperature to block non-specific binding sites. Color was developed using the DAB Substrate Kit (Zsbio, ZLI-9017). The primary antibodies used in this study included anti-human TMEM132A antibody (1:200, Merck, ABC148), anti-human perilipin-1 (1:500, Immunoway, PT0458R), and anti-human perilipin-2 (1:200, Abclonal, A6276). Slides were examined and images were captured using a laser scanning microscope (Zeiss, Axioscan 7).

Multiplex IHC (mIHC) was performed using the Opal 4-Color Manual IHC Kit (Akoya Biosciences, NEL810001KT) according to the manufacturer’s instructions. Briefly, following dewaxing in xylene and rehydration through a graded ethanol series, FFPE tissues were fixed with 10% neutral-buffered formalin. Antigen retrieval was carried out with Opal AR6 Buffer using high-pressure heating. After cooling and blocking, sequential staining with primary antibodies, HRP-conjugated polymers, and Opal fluorophores was performed, and the cycles were repeated until all markers had been stained. The primary antibodies used in this study included anti-human TMEM132A (1:200, Merck, ABC148), anti-human SREBP1 (1:500, Huabio, HA722160), anti-human SREBP2 (1:200, Abclonal, A13049), anti-human EGFR (1:200, Huabio, ET1701-22), anti-human Rab7 (1:1000, Huabio, ET1611-96), and anti-human Rab11A/B (1:200, Abclonal, A20996). Finally, nuclei were counterstained with DAPI. Slides were examined and images were captured using a confocal microscope (Zeiss, LSM900).

### Biomolecular interaction prediction with AlphaFold

The full-length amino acid sequences of TMEM132A (UniProt ID: Q24JP5) and EGFR (UniProt ID: P00533) were used as input for protein complex structure prediction, which was performed using AlphaFold3 via the online official AlphaFold Server (https://alphafoldserver.com)^55^. Five independent predictions were generated with default parameters (recycles = 3, ensemble mode enabled). The output model with the highest interface predicted template modeling (ipTM) scores was selected for downstream analysis. The predicted structure was drawn using PyMOL (v3.0)^56^.

### TCGA data analysis

For TCGA survival analysis of TMEM132A and EGFR co-expression, clinical and RNA-seq data for the TCGA-KIRC, TCGA-LUAD, and TCGA-BRCA cohorts were downloaded from the Genomic Data Commons (GDC) using the TCGAbiolinks R package. Clinical data in BCR XML format were parsed using the XML package, and samples with incomplete or unpartable clinical records were excluded. Only primary tumor samples were retained for downstream analysis. Gene expression matrices were extracted from RangedSummarizedExperiment objects using the tpm_unstrand assay. Ensembl gene identifiers were stripped of version numbers and converted to HGNC gene symbols using biomaRt. Duplicated genes were collapsed by summing expression values, and genes with mean TPM≤5 were removed. Clinical and expression data were integrated by patient barcode. Overall survival time was defined as the maximum of days_to_last_known_alive, days_to_death, and days_to_last_followup, and survival status was coded as 1 for deceased patients and 0 for living patients. Within each cancer type, TMEM132A and EGFR expression levels were dichotomized independently according to the cohort-specific median. Following the co-expression stratification logic reported by Yu et al^57^, patients with both TMEM132A and EGFR expression above the median were assigned to the “both high” group, whereas all remaining patients were assigned to the“single or simultaneous low” group. Kaplan–Meier overall survival curves were generated using the survival and survminer R packages, and statistical significance was assessed by the log-rank test.

Protein-level correlation between TMEM132A and EGFR in breast cancer was examined using the cBioPortal web platform. Protein data were queried through cBioPortal from the Breast Invasive Carcinoma (TCGA, PanCancer Atlas) study, using the mass spectrometry by CPTAC protein profile available for that cohort.

### Nanobody Screening

The screening of nanobodies was performed by Immunoway Biotechnology Co., Ltd. In brief, three poly-peptide antigens (CWWRRLRASLR, CAPLLPLRIELT, CVDFWWRRLRASL) were synthesized based on the interaction sites between the two proteins. Mixture of the three peptides were incubated with the CamNano phage display library (containing hundreds of billions of clones). Six rounds of biopanning with the peptide antigens were performed, followed by limiting dilution or single-clone isolation to select monoclonal phages. Recombinant nanobody crude extracts were then obtained. individual extract was subjected to ELISA validation for binding efficiency to the fixed antigen mixtures. The following criteria: positive well reading / negative well reading ≥3 and positive well reading - negative well reading ≥1 was used for screening positive clones. Positive clones were sequenced to determine the amino acid sequence of the nanobodies.

### Expression and Purification of Nanobodies

The eukaryotic expression and purification of nanobodies were performed by Immunoway Biotechnology Co., Ltd. The coding sequence of the nanobody was fused with the mouse IgG1 Fc region, placed downstream of the IL-2 signal peptide sequence, and cloned into the pcDNA3.4 vector under the CMV promoter. The recombinant plasmid was transfected into HEK293F cells using polyethylenimine (PEI) Max (Polysciences, 24765-1). The cell supernatant was centrifuged (1,500 rpm, 10 min), filtered through 0.22 μm or 0.45 μm membranes, and purified via Protein G affinity chromatography to obtain the nanobody-Fc fusion protein. The purity of the purified protein was verified by SDS-PAGE.

### *In vivo* study of anti-tumor effect of LFNanoT132A#1 and #3

Syngeneic model from mice 4T1 and panc02 cells were exployed to evaluate the *in vivo* anticancer efficacy of LFNanoT132A#1 and #3. When the tumor reached 50–100 mm^3^, mice were randomly assigned into different treatment groups. Control solution (PBS) and LFNanoT132A#1 and #3 were injected into tumors at a dose of 10mg/kg body weight each two days intravenously. When the tumor reached 100-200 mm^3^, the mice were divided randomly into different groups. PBS, LFNanoT132A#1 or #3 were injected into mice at a dose of 50μg/mice three times every week intratumorally. Body weights and tumor volumes of mice were measured prior to each administration until the endpoint. The end point for maximum tumor larger diameter was 2cm.

### *In vivo* toxicity of LFNanoT132A

For intratumoral injection, mice were euthanized after six administrations. Major organs (heart, liver, spleen and kidneys) were harvested for histopathological examination. For intravenous injection via the tail vein, after five administrations, the same measurements were performed.

### Statistical analysis

All the experiments were performed at least three times independently. All the data were presented as mean ± SEM. Statistical analyses were performed using graphpad software (version 8.0.1.244), for comparison between two cell groups, p values were analysed using two-tailed student’s t-test unless otherwise indicated. For multiple comparisons, analysis of variance (ANOVA) was performed. For tumour volume, statistical significance was examined through two-way ANOVA analysis. To compare data between two animal or patient samples, Mann-Whitney test was used. *p<0.05, **p<0.01, ***p<0.001 was considered as statistically significant; ns, no significance. The correlation coeffients were determined using pearon’s rank correlation test.

## Acknowledgement

We are particularly grateful to Dr. Ke Shuai and Sumin Feng, the committee members of this project, for their assistance and valuable comments on this work. This work was supported in part by the National Natural Science Foundation of China (grant 82573830 to Xijuan Liu, grant 82472787 to Jun Wang, grant 82204454 to Wentong Fang); the Fundamental Research Funds for the Central Universities (KG202504 to Xijuan Liu and Jun Wang).

## Author contributions

XJ.L. (Xijuan Liu), J.W (Jun Wang), YR.F. (Yaru Fu), XY. W (Xiyi Wei) and C.Q. (Chao Qin) designed, performed and interpreted the experiments and co-wrote the paper. YR.F. also contributed to cell phenotype, Co-IP, IFA, IP-MS and RNA-sequencing assays; QQ.N. (Qiqi Ni), C.N. (Can Ning) and JH.W.(Junhao Wang) helped with IHC, mIHC and mice experiments; QQ.N., C.Z. (Cheng Zhang) and MY.W.(Mingyu Wu) contributed to PDOs and mice mating and breeding; X.F.(Xiang Fang) helped biomolecular interaction prediction with AlphaFold; JX.W.(Jixian Wang) and JH.W. helped with Bioinformatic analysis and TCGA data analysis; JY.Q.(Jiayi Qian), WT.F. (Wentong Fang), L.G. (Li Gong) and J.Y. (Jing Yao) provided clinical samples for PDOs; D.Z. (Dan Zhang) and XM.L. (Xiaoming Li) provided clinical samples for IHC and mIHC; F.Z. (Fei Zhao) provided cancer cell lines and helped with manuscript editing; NH.S. (Ninghong Song) provided ccRCC TMAs; YQ.H.(Yuanqiao He) helped with mice assays with LFNanoT132A#1 and #3.

---

**Figure S1 TMEM132A regulates ccRCC cell proliferation, anchorage-independent growth.**

(**A**) Schematic overview of the CRISPR-Cas9 screening performed in 786-O Cas9 stable cell lines. (**B**) ChIP-seq binding peaks of SFMBT1 and ZHX2 on the TMEM132A locus (upper); Venn diagram showing the overlap of downstream target genes of SFMBT1 and ZHX2 in ccRCC (lower); Data were derived from GSE141577 (SFMBT1) and GSE109953 (ZHX2), respectively. (**C-D**) qRT-PCR quantification (**C**) and immunoblots (**D**) to detect TMEM132A level from indicated ccRCC paired patient tumor (T) tissues and adjacent non-tumor (N) tissues. (**E**) qRT-PCR quantification (upper) and immunoblots (lower) to detect TMEM132A level from indicated cell lines. (**F-G**) Immunoblots for lysates (**F**) and 3-D soft agar assays (**G**) of indicated ccRCC and HKC cells with TMEM132A depletion. (**H-K**) Immunoblots for lysates (**H**), CCK8 assays (**I**), colony formation assay (upper) and 3-D soft agar assays (lower) (**J**), and quantification of soft agar assays (**K**) of 786-O cell lines cells with lentivirus encoding EV or Res132A, followed by TMEM132A depletion. *sh2 versus shCtrl; ^#^sh2+Res132A versus sh2; ^&^shCtrl+Res 132A versus shCtrl. (**L-O**) Immunoblots for lysates (**L**), CCK8 assays (**M**), colony formation assay (upper) and 3-D soft agar assays (lower) (**N**), and quantification of soft agar assays (**O**) of 769-P cells transduced with lentivirus expressing either sgRNA control (sgCtrl) or TMEM132A CRISPR sgRNA (sg1 and sg2). *sg2 versus sgCtrl; ^#^sg1 versus sgCtrl. (**P-S**) Immunoblots for lysates (**P**), CCK8 assays (**Q**), colony formation assay (upper) and 3-D soft agar assays (lower). (**R**) and quantification of soft agar assays (**S**) of 786-O cells transduced with lentivirus expressing either inducible shCtrl or TMEM132A sh2/5 with DOX treatment. *sh5 tet-on versus shCtrl tet-on; ^#^sh2 tet-on versus shCtrl tet-on. Data are shown as mean ± SEM, Statistical significance was determined by one-way ANOVA (**C**, **E**, **K**, **O** and **S**) and two-way ANOVA (**I, M** and **Q**).

**Figure S2. Knockout of TMEM132A Drastically Suppressed Tumor Growth**. (**A**) HR values of TMEM132 family members across pan-cancer type. (**B-C**) Immunoblots for lysates (**B**), CCK8 assay (**C**, left), 3-D soft agar assays (**C**, middle) and quantification of soft agar assays (**C**, right) of indicated cells with TMEM132A depletion. (**D**) Representative hematoxylin and eosin (H&E) staining of TNBC organoid. (**E**) Representative H&E staining of tumor tissue from mice after orthotopically injection with TNBC PDX cells. (**F-H**) Immunoblots for lysates (**F**), 3-D soft agar assays (**G**) and quantification of soft agar assays (**H**) of EMT6 cells with TMEM132A depletion. Data are presented as means ± SEM. Statistical significance was determined by unpaired Student’s *t* test.

**Figure S3. TMEM132A Modulates Acetyl-CoA Production by Transcriptionally Regulating the Key Synthetases ACLY and ACSS2.** (**A**) Volcano plot showing differentially expressed genes (DEGs) from RNA-seq of 786-O cells (TMEM132A sh5 vs. shCtrl). (**B**) Unbiased hierarchical clustering was performed to visualize significantly changed lipid-related metabolites (P < 0.05) in 786-O cells following TMEM132A knockdown by shRNAs. (**C**) 3-D soft agar assays of indicated cells with TMEM132A depletion followed DCA treatment. (**D**) Representative IHC staining of PLIN1 and PLIN2 in BRCA patient tissues stratified by high (TMEM132A^high^) or low (TMEM132A^low^) TMEM132A expression. (**E**) QRT-PCR quantification of *ACSS2*, *ACLY* and *TMEM132A* level in the indicated cell lines with lentivirus encoding empty vector (EV) or Res132A, followed by TMEM132A depletion. Statistical significance was determined by two-way ANOVA. *sh5. versus shCtrl; ^#^sh5+ Res132A versus sh5. (**F**) Immunoblots for lysates from xenograft tumors derived from mice injected with indicated cells with TMEM132A depletion. (**G**-**H**) 3D soft agar assays (**G**) and quantification (**H**) of 3D soft agar assays of 786-O cells with TMEM132A OE followed with or without ACLY inhibitor and/or ACSS2 inhibitor treatment as indicated concentration. Statistical significance was determined by one-way ANOVA. *TMEM132A OE versus EV; ^#^ACLYi/ACSSi/ACLYi+ACSSi versus TMEM132A OE. Error bars represent SEM.

**Figure S4. TMEM132A post-translationally regulates EGFR protein stability.** (**A**) Partial list of FLAG-TMEM132A interactors identified by FLAG IP-MS in 786-O cells. (**B**) Immunoprecipitations and immunoblots for lysates from EV and TMEM132A OE 786-O cells or 293T cells. (**C**) Kaplan–Meier plot of overall survival of patients from the TCGA-KIRC, TCGA-LUAD, and TCGA-BRCA cohorts stratified by TMEM132A and EGFR co-expression level. Risk tables are shown below each plot. (**D**) Immunoblots for indicated cell lysates with TMEM132A OE (left) or depletion (right). (**E**) Immunoblots for indicated cell lysate with TMEM132A OE (left) or depletion (right) followed CQ treatment. (**F**) Ubiquitination level of endogenous EGFR in indicated cell lines following TMEM132A OE (left) or depletion (right). (**G**-**H**) Immunoblots for indicated cell lysates with TMEM132A OE (**G**) or depletion (**H**) followed with 100μM CHX treatment at indicated intervals. The right panel shows the quantification of EGFR bands, with tubulin used as the loading control. EGFR expression levels in the EV/OE and shCtrl/sh5 groups at the 0-hour time point were normalized to 1.0. (**I**) Immunoprecipitations and immunoblots for lysates from MDA-MB-231 cell lines with TMEM132A OE. (**J-K**) Representative IFA images showing co-localization of EGFR and Rab7 or Rab11A in indicated cell lines with TMEM132A OE (**J**) or depletion (**K**). Quantitative analysis of co-localization was shown as Pearson’s coefficient. (**L-M**) Immunoblots for cell lysates (**L**), 3-D soft agar assays (**M**) in indicated cell lines transduced with lentivirus encoding EV or EGFR followed by TMEM132A depletion. Error bars represent SEM. Statistical significance was determined by unpaired Student’s *t* test (**J** and **K**).

**Figure S5. The TMEM132A-EGFR Axis Promotes SREBP Nuclear Translocation.** (**A**) Relative mRNA levels of SREBP1 or SREBP2 analyzed by RNA-seq in 786-O cells with TMEM132A depletion. (**B-C**) Immunoblots analysis for the indicated proteins with TMEM132A OE (**B**) or depletion (**C**) with or without EGF (10 ng /ml) treatment for 1 h. (**D-E**) MDA-MB-231 and A549 cells with TMEM132A OE (**D**) or depletion (**E**) were subjected to fractionation, followed by immunoblots analysis for the indicated proteins. (**F**-**G**) Representative IFA staining of SREBP1 or SREBP2 in indicated cells with TMEM132A OE (**F**) or depletion (**G**).

**Figure S6.** (**A**) Alpha-fold3 predicted interaction interface between TMEM132A and EGFR. Dashed lines indicate hydrogen bonds between amino acid residues of TMEM132A and EGFR. (**B**) Immunoprecipitations and immunoblots of 293T cells transfected with plasmids encoding EV or FLAG-TMEM132A (aa392-635). (**C**) Immunoprecipitations and immunoblots of 769-P cells transduced with lentivirus encoding EV or FLAG-TMEM132A (aa392-635). (**D**) Immunoprecipitations and immunoblots of 293T cells transfected with plasmids encoding EV or EGFR (aa25-645). (**E**) Immunoprecipitations and immunoblots of 293T cells transfected with indicated plasmids. (**F**) Immunoblot analysis of cell lysates from 293T cells transfected with the indicated plasmids expressing WT or mutant-TMEM132A. (**G**) Immunoprecipitations and immunoblots of 293T cells transfected with the indicated plasmids expressing WT or mutant-TMEM132A with CQ treatment for 24h. (**H**) Immunoblot analysis of cell lysates from 769-P cells (left) and UMRC-2 cells (right) transduced with lentivirus encoding WT or mutant-TMEM132A. (**I**-**J**) 3-D soft agar assays (left) and quantification of soft agar assays (right) of 769-P cells (**I**) and UMRC-2 cells (**J**) transduced with lentivirus encoding WT or mutant-TMEM132A. (**K**-**N)** 3-D soft agar assays (left) and quantification (right) of 3-D soft agar assays of SK-N-BE2 (**K**), LNCaP-C4-2 (**L**), BxPC3 (**M**) and A549 (**N**) cells with 10μg/ml of LFNanoT132A#1 or #3 treatment. Error bars represent SEM. Statistical significance was determined by one-way ANOVA.

**Figure S7.** (**A-C**) Tumor images (**A**), tumor volume (**B**), body weight (**C**) of Balb/c mice implanted with 4T1 tumor tissue cubes and then intratumorally injected with LFNano132A#1 or #3. (**D**) Representative microscopic examination of H&E-stained tissue sections of the indicated organs of Balb/c mice implanted with 4T1 tumor tissue cubes and then intratumorally injected with LFNano132A#1 or #3. (**E**) Representative microscopic examination of H&E-stained tissue sections of the indicated organs of Balb/c mice implanted with 4T1 tumor tissue cubes and then intravenously injected with LFNano132A #3. Data are presented as means ± SEM. Statistical significance was determined by Mann-Whitney test.

