## Supplementary figures and images for "Dissecting the TMEM132A-EGFR Dependency to Unlock Translational Therapeutic Opportunities for Pan-Solid Tumor"

### Figure S1

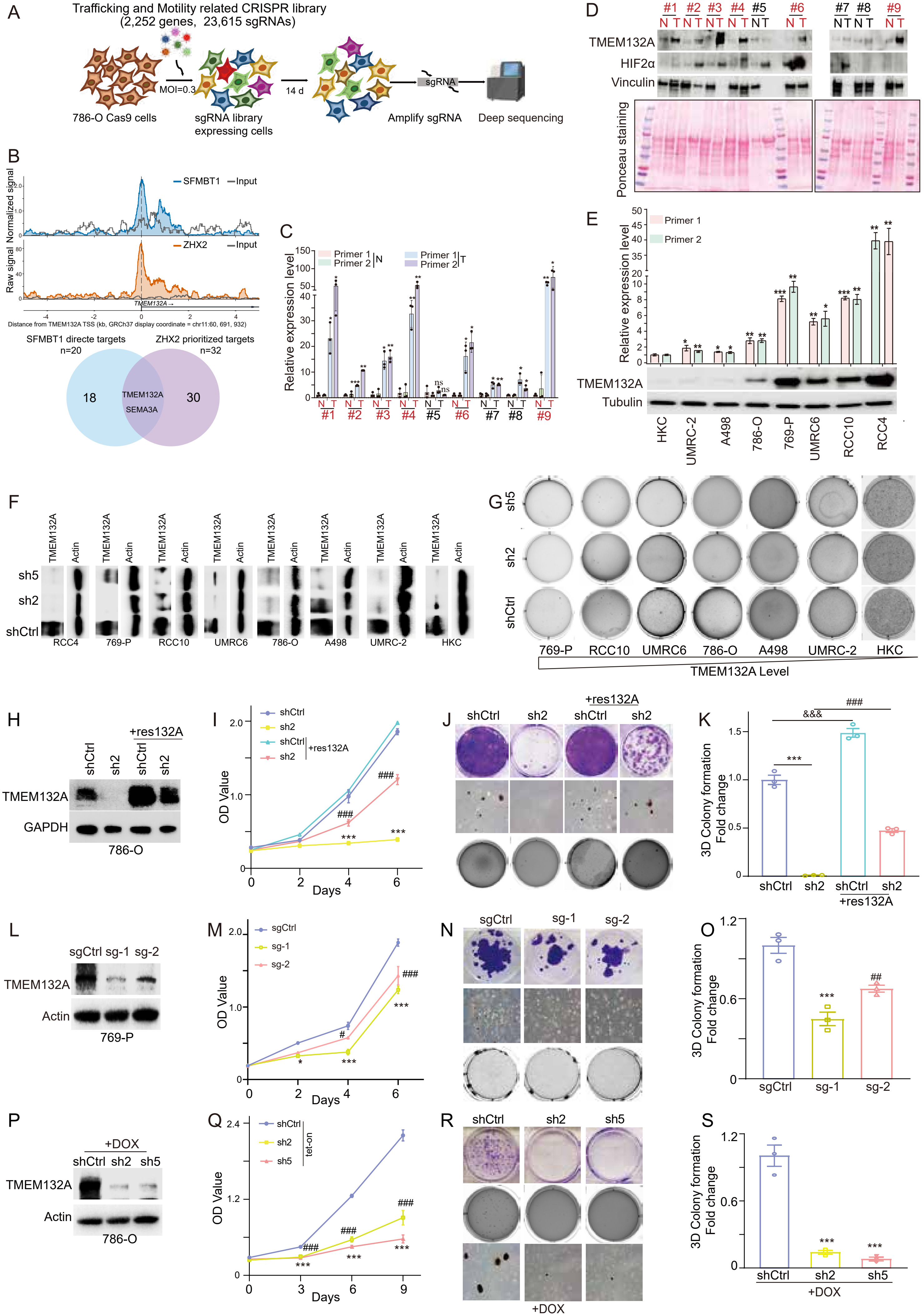

### Figure S2

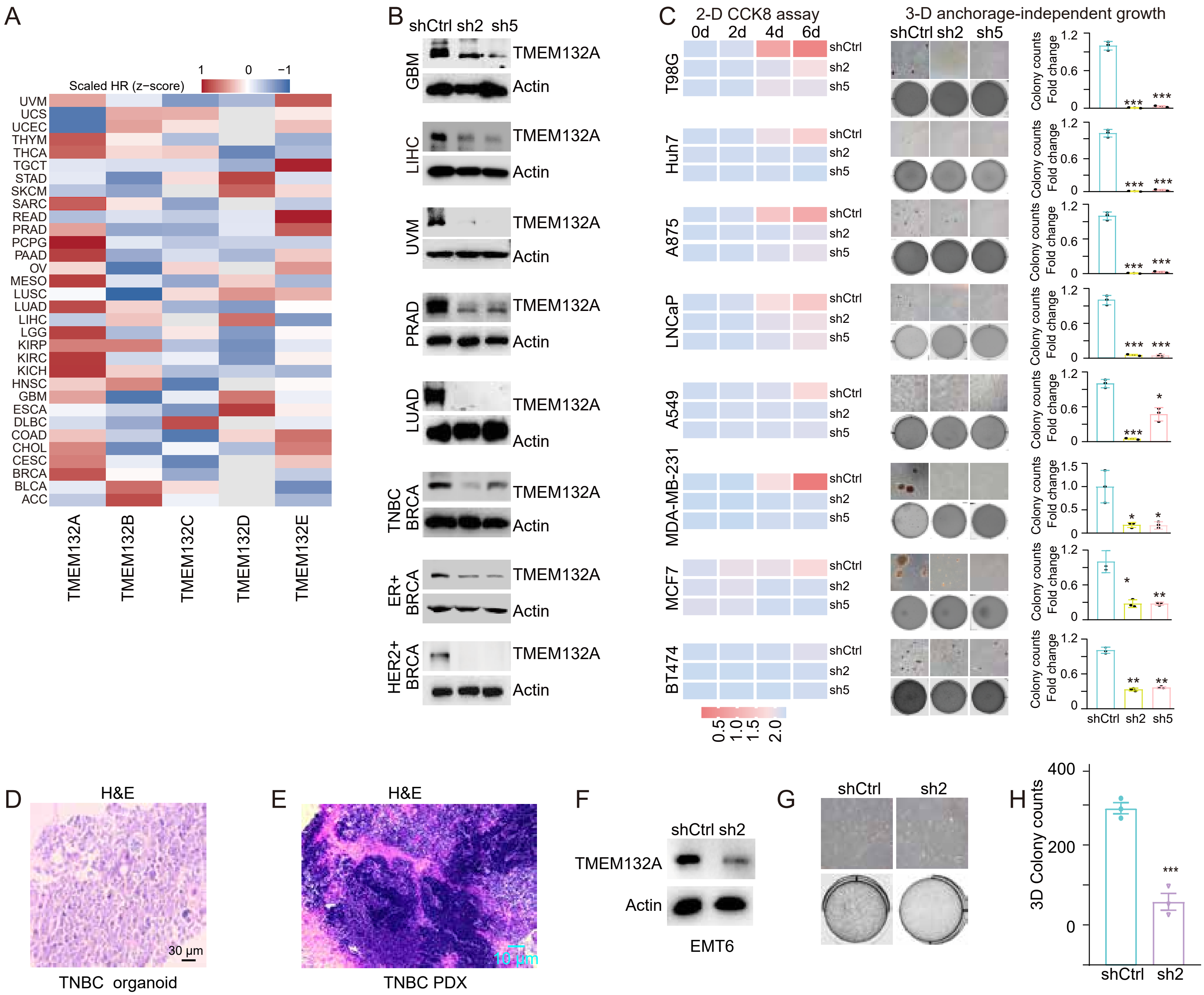

### Figure S3

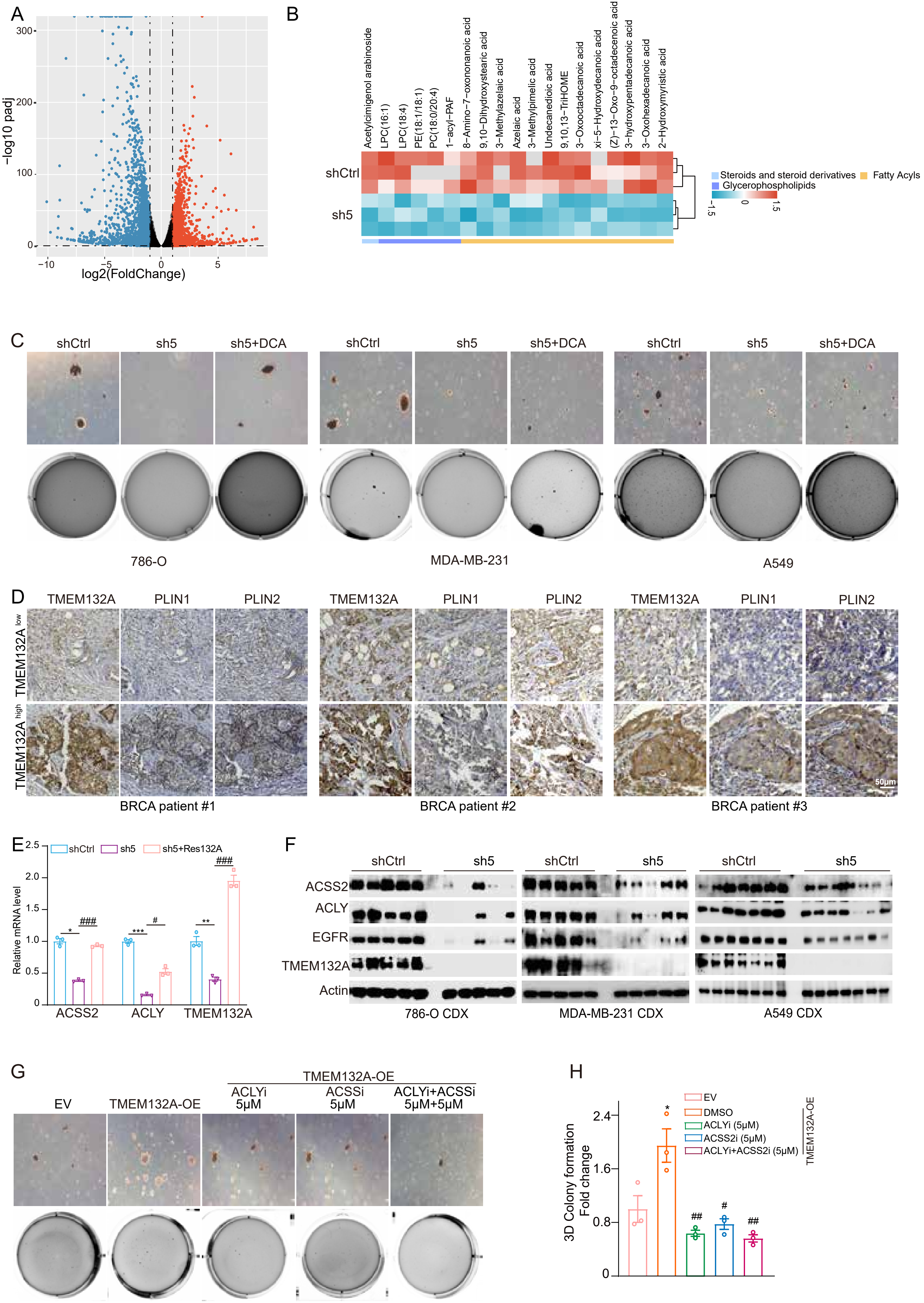

### Figure S4

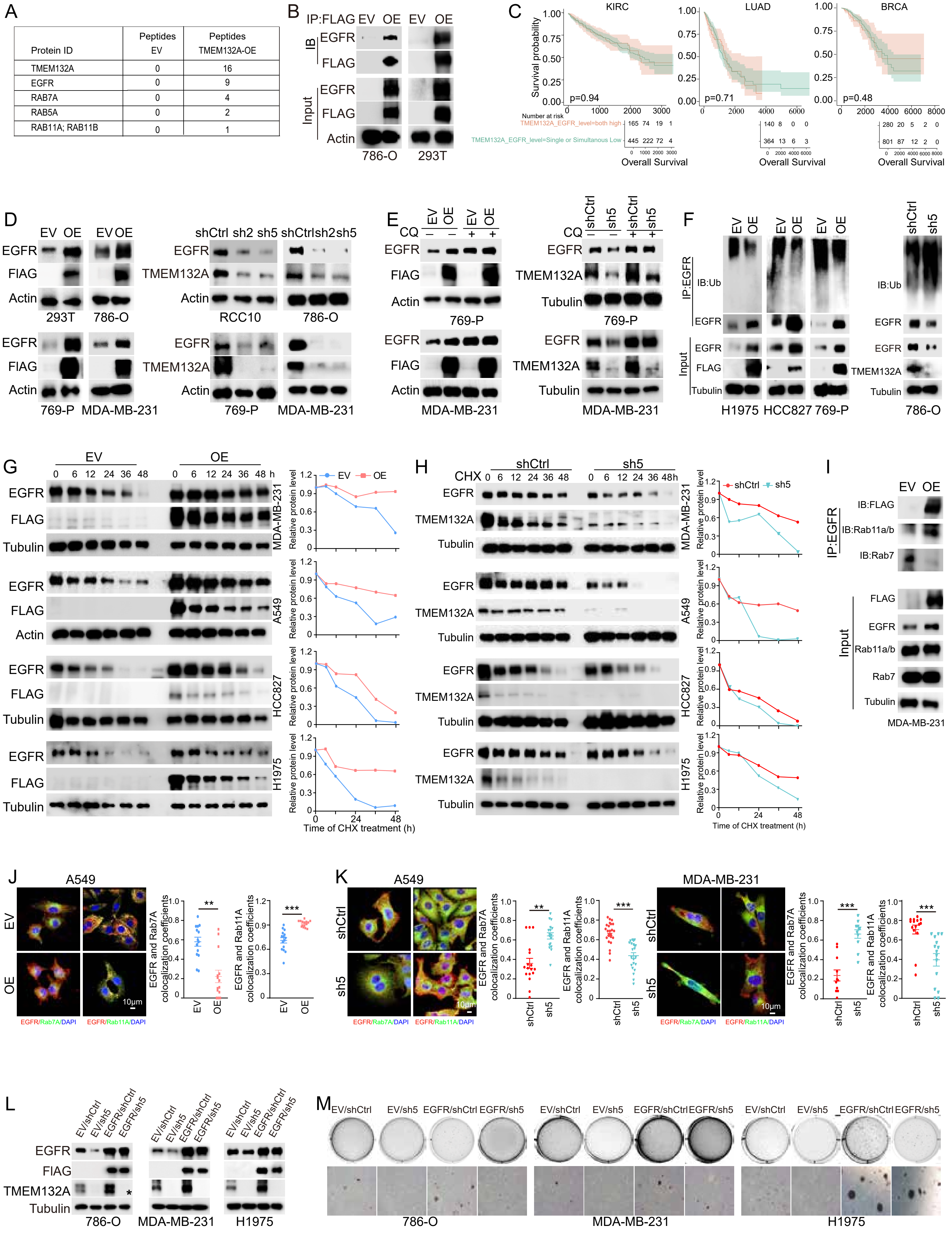

### Figure S5

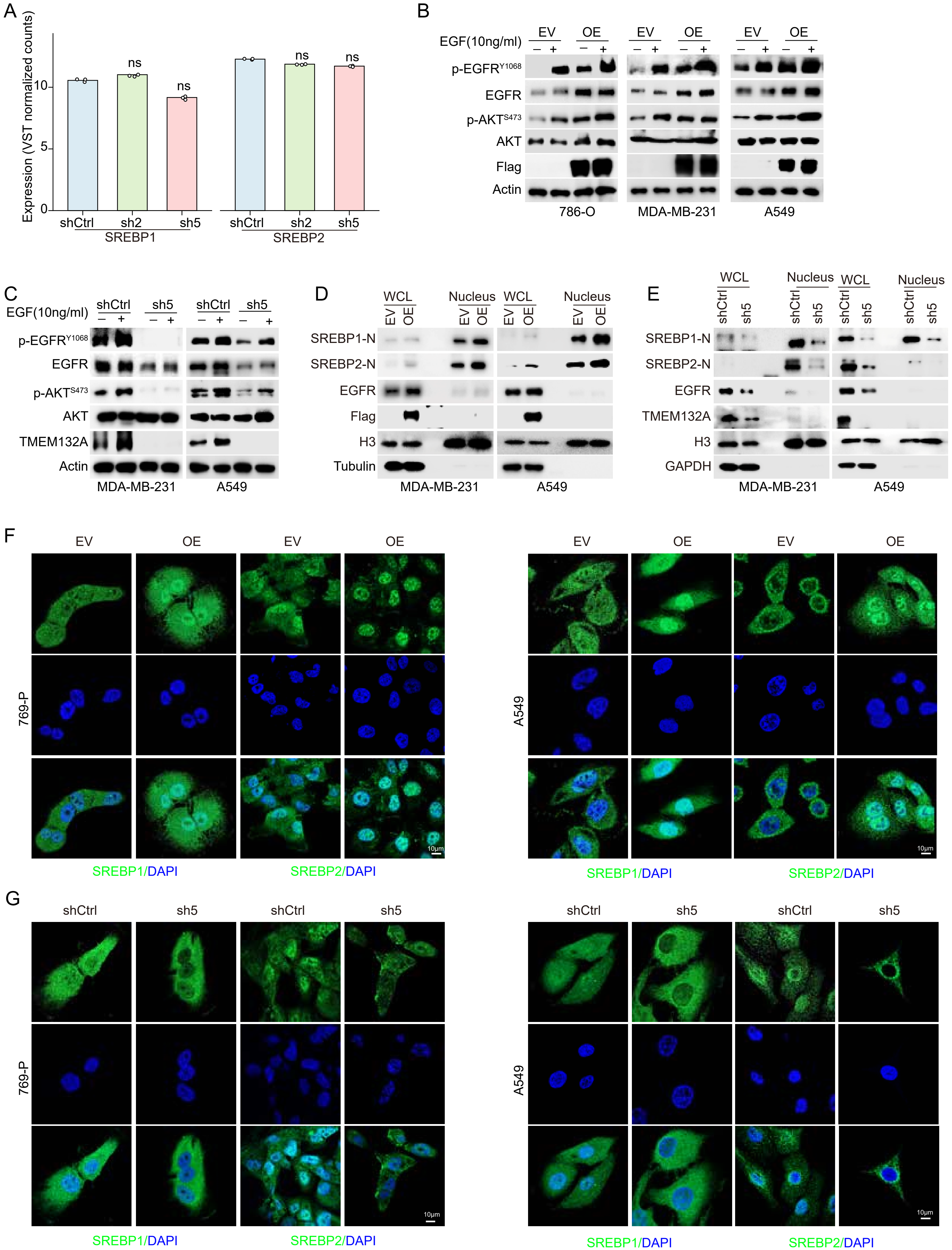

### Figure s6

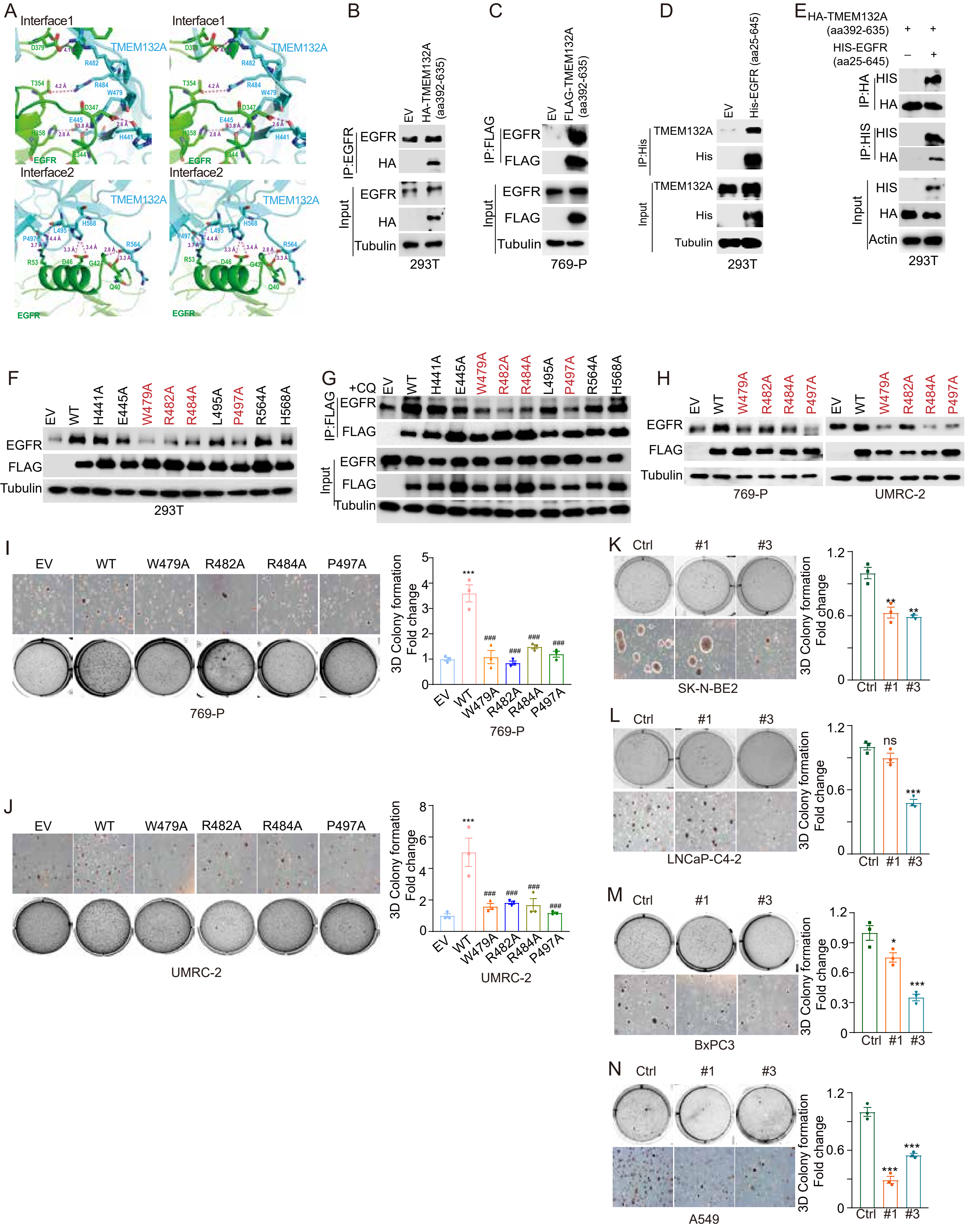

### Figure S7

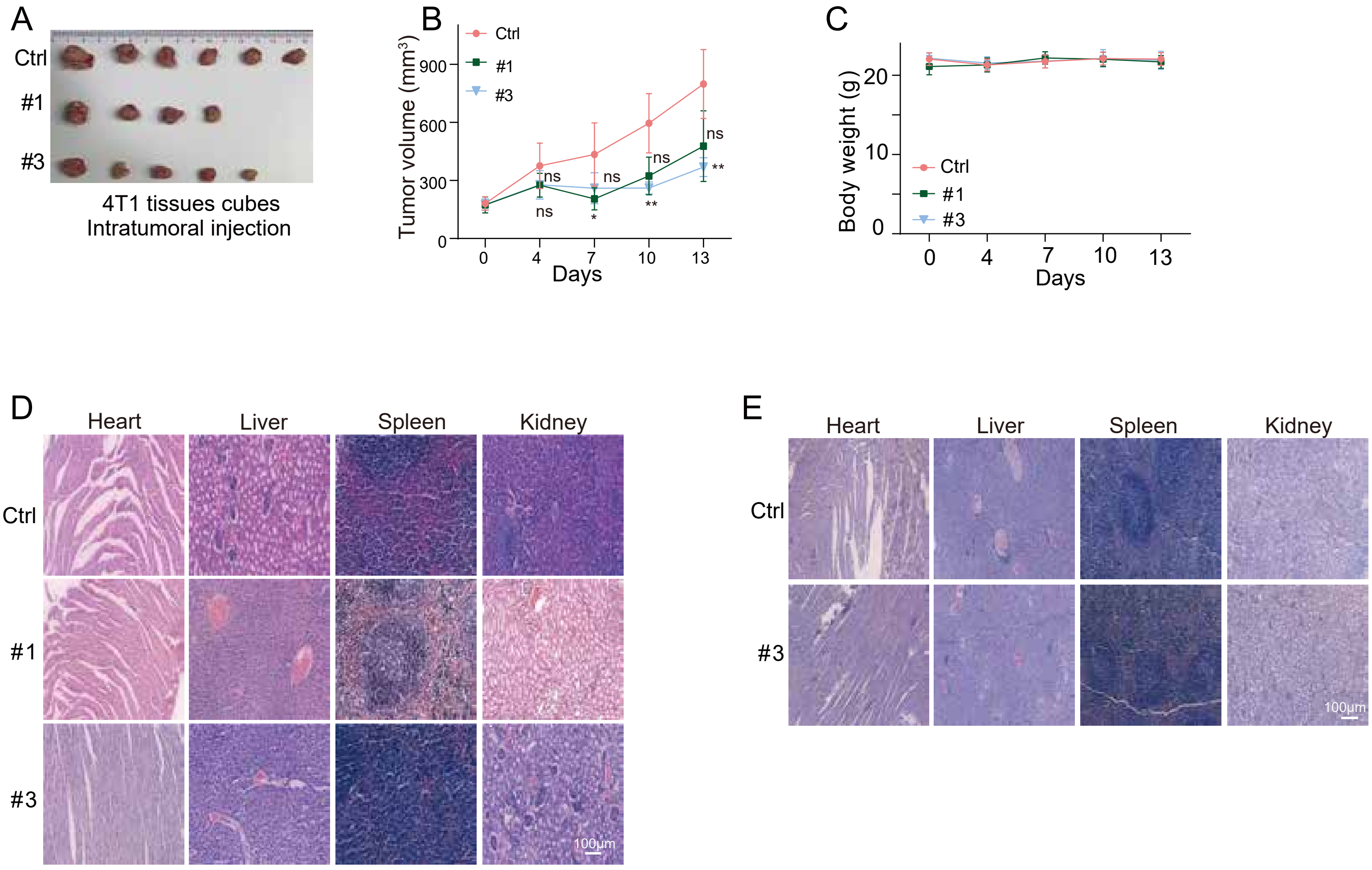

### Supplementary data

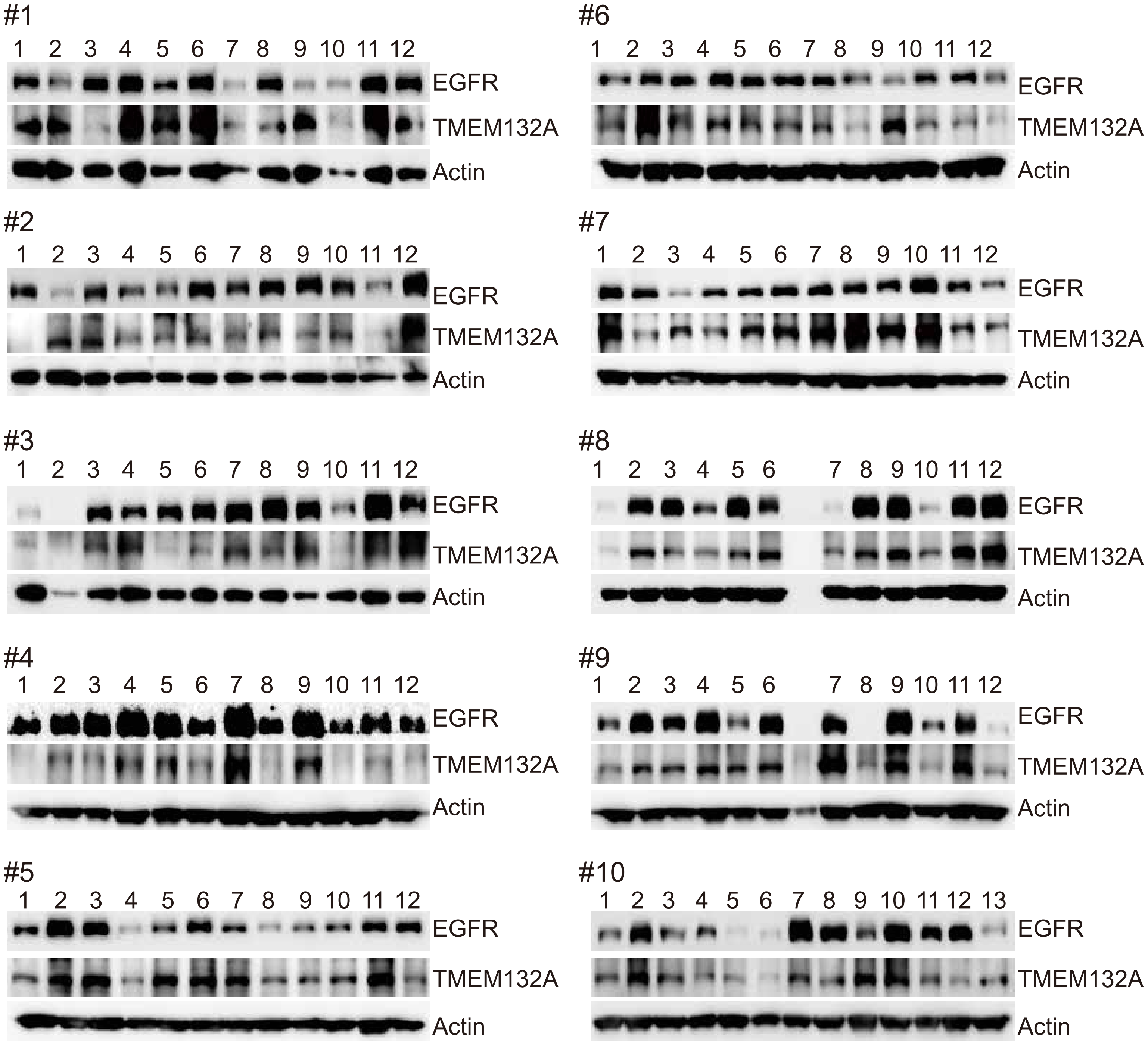
